# Decoding silence in free recall

**DOI:** 10.1101/2021.01.26.428351

**Authors:** Francesco Fumarola, Zhengqi He, Łukasz Kuśmierz, Taro Toyoizumi

## Abstract

In experiments on free recall from lists of items, not all memory retrievals are necessarily reported. Previous studies investigated unreported retrievals by attempting to induce their externalization. We show that, without any intervention, their statistics may be directly estimated through a model-free analysis of inter-response times – the silent intervals between recalls. A delay attributable to unreported recalls emerges in three situations: if the final item was already recalled (“silent recency effect”); if the item that, within the list, follows the latest recalled item was already recalled (“silent contiguity effect”); and in sequential recalls within highly performing trials (“sequential slowdown”). We then turn to reproducing all these effects by a minimal model where the discarding of memories (“bouncing”) occurs either if they are repetitious or, in strategically organized trials, if they are not sequential. Based on our findings, we propose various approaches to further probing the submerged dynamics of memory retrieval.

## Introduction

Recall of memories in humans is increasingly regarded as a multi-layered process emerging from the complex interplay of free association and higher cognitive functions (Becker & Lim, 2003; Gronlund & Shiffrin, 1986; Norman, Yeagle, Harel, Mehta, & Malach, 2017; Polyn, Norman, & Kahana, 2009; Raaijmakers & Shiffrin, 1981). Various neuroimaging techniques are being deployed to study what has been termed *post-retrieval* processes, a set of processes aimed at monitoring and evaluating each retrieved memory (Badre, Poldrack, Paré-Blagoev, Insler, & Wagner, 2005; Hayama, Johnson, & Rugg, 2008; Hayama & Rugg, 2009; Roediger III & Tulving, 1979; Rugg, Johnson, & Uncapher, 2015). Evidence suggests a wide participation in the top-down control of memory recall by prefrontal (Long, Öztekin, & Badre, 2010; Savage et al., 2001), frontal (Stuss et al., 1994), and parietal (Cabeza, Ciaramelli, Olson, & Moscovitch, 2008) areas. Ventrolateral prefrontal cortex (VLPFC), in particular, has been hypothesized to support a post-retrieval selection mechanism (Badre et al., 2005) with selection demands increasing when task-irrelevant representations are prepotent; electrodes implanted in VLPFC (Norman et al., 2017) were found to increase their activity a few hundred milliseconds after the onset of the recall stage, consistent with a top-down post-retrieval feedback. The general breadth and organization of post-retrieval processes, however, remains unknown.

An ideal setting for addressing the question is the experimental paradigm known as *free recall* (Bousfield & Bousfield, 1966; Ebbinghaus, 1913; Murdock Jr, 1962; Postman & Phillips, 1965; Tulving, 1962), which has been providing insight on all facets of human episodic memory for over a century (Binet & Henri, 1894; Kirkpatrick, 1894). Free-recall experiments are split into a presentation stage and a recall stage; the former consists in the presentation of a list of items (often words), the latter in the recall of those items (usually reported verbally by subjects) with no constraint on the ordering. The aim is to maximize performance, defined as the number of distinct items recalled from the list. (For a comprehensive monograph on both experiments and theories see Kahana (2012)).

Results on free recall accumulated in the literature may indeed be subsumed under two classes: results whose broad understanding does not seem to require invoking post-retrieval mechanisms; and results that are, more or less implicitly, underpinned by the existence of post-retrieval activity.

Among the former group, the most studied effects concern the *serial position* of recalled items, i.e. their order within the list presented. These include the *recency* and *primacy* effects (Murdock Jr, 1962), i.e., the tendency to more consistently recall items respectively from the beginning and end of the list. The difference in serial position between two consecutively recalled items is known as “serial-position lag”, and the *lag-recency* or *contiguity* effect (Kahana, 1996) is a bias toward small lags, i.e. the tendency to recall contiguously items that are contiguous within the presented list (Healey, Long, & Kahana, 2019; Howard & Kahana, 1999). The additional tendency to recall in forward order (“forward asymmetry”) makes lag *L* = +1 the most likely transition, which we will refer to as “sequential” (Kahana & Caplan, 2002).

In parallel, there is a body of work showing evidence for postretrieval mechanisms such as, notably, repetition avoidance. Repetition avoidance is indirectly evidenced by various experimental facts. For example in serial recall, where items must be recalled in their order, fewer intrusions were found to correlated with better recall of the final item (Farrell & Lewandowsky, 2012); this was attributed to the fact that recall of the final item is boosted by repetition avoidance and becomes harder if fewer items are blocked as repetitious. More detailed descriptions of repetition avoidance were worked out with other models (Lohnas, Polyn, & Kahana, 2015; Vousden & Brown, 1998). Since it is difficult to examine all possible models, it would be desirable to develop model-free approaches to extract information on unreported events from ordinary free-recall data.

Another topic that has been the focus of a large amount of attention is the observed adoption of various retrieval strategies such as chunking, *i.e.* retrieving consecutive items in one group or serializing, *i.e.* retrieving items in their serial order (Becker & Lim, 2003; Gianutsos, 1976; Gronlund & Shiffrin, 1986; Romani, Katkov, & Tsodyks, 2016). Such strategies are mostly consistent within a trial and considered to be a characteristic of trials rather than recalls. They were shown to be effective in boosting performance but are implemented by a minority of subjects and even by those only in a fraction of trials (Romani et al., 2016). In fact, they are often developed and refined over the length of an experimental session as each subject learns from his performance in previous trials (Keppel, Postman, & Zavortink, 1968). While some recall strategies are related to chunking, it is long since known that a particularly advantageous strategy consists in waiving the *freedom* perk of free recall by serializing (Gianutsos, 1976; Romani et al., 2016; Zhang, Griffiths, & Norman, 2021). Sticking to such strategies should involve discarding any retrieval that does not fit the prescribed recall sequence.

One approach to unraveling the histories of unreported recalls consists in directly demanding subjects to externalize them. This type of interventionalism was attempted as early as with the “unedited recall” experiments of Hogan (1975), where subjects were instructed to express all items that came across their mind, and has provided further support to the idea that termination is predominantly triggered by repetition (Unsworth, Brewer, & Spillers, 2010). Although such procedure was shown to yield results coherent with the “edited” (i.e. regular) version of free recall (Roediger & Payne, 1985) and was consequently proposed as a method to explore age dependence (Kahana, Dolan, Sauder, & Wingfield, 2005), memory capacity (Unsworth & Brewer, 2010), and intrusions (Lohnas et al., 2015), self-reporting obviously involves physical bounds on the time resolution and, potentially, biases created by the amount of instruction (Aguirre, Gómez-Ariza, & Bajo, 2020). Whether or not such data offer an exact representation of thought processes taking place during ordinary free recall is a question that only electrophysiological inquiry could address in full.

Here, we sidestep both the caveats of the externalization paradigm and, in our opening foray, the problem of selecting a model. We focus instead on the model-free analysis of one specific behavioral observable – the time that elapses between consecutive recalls, also known as inter-recall interval or IRI (Healey, Crutchley, & Kahana, 2014; Patterson, Meltzer, & Mandler, 1971; Pollio, Kasschau, & Denise, 1968; Pollio, Richards, & Lucas, 1969).

During the recall stage of a typical free-recall trial, the first few recalled items are reported rather quickly; in contrast, towards the end of the trial the IRIs drastically increase (Murdock & Okada, 1970; Rohrer & Wixted, 1994). Many features of this growth were shown to be reproduced by a pure-death process, in which items are sampled with replacement from a pool of listed items (McGill, 1963; Rohrer & Wixted, 1994; Vorberg & Ulrich, 1987). Crucially, in that model it is assumed that only items that have not been reported so far are admissible, and thus, when the other items are drawn, they are simply discarded. Because towards the end of a trial there are many previously recalled (hence non-admissible) items, many samples have to be drawn on average and thus IRIs increase. Although the pure-death model is very simplistic, its structure suggests that a form of post-retrieval editing takes place during the recall phase and that only some of the sampled (retrieved) items are reported.

We will argue that a careful but straightforward analysis of both the history of reported recalls and the recorded IRIs can allow to uncover hidden retrievals that are suppressed by the participants. Most of the inferred instances of discarded retrievals are related to the avoidance of repetitions of already-recalled items, but our analysis strongly suggests that some of the participants suppress even novel recalled items in order to adhere to their retrieval strategy.

In the second part of the paper, we will buttress our model-free analysis of data by means of a toy model chosen for its simplicity and capability to explain simultaneously different types of unreported recalls.

## Results

### Two free recall phenomena

The data we considered, collected by the Computational Memory Lab at the university of Pennsylvania, concern experiments performed with lists of *N* = 16 words (see Materials and Methods for details). The dataset displayed standard serial-position effects (with recency and primacy emerging as in Fig. 10).

We begin by pointing out two general facts in this dataset:

a. Although repetitions are not expressly forbidden and are duly recorded by the experimenter, the number of repetition-free trials is about twice the chance level set by a fitted Markovian transition model (1A) of the type first proposed in Hogan (1975). This fact agrees with previous inquiries (see Introduction) and indicates that subjects have a spontaneous tendency toward not reporting twice the same recall (see Fig. 11 for a more detailed comparison of the repetition statistics to chance level in the dataset, and see its caption for a description of the Markovian transition model).
b. We define “trial sequentiality” as the fraction of sequential recalls in the trial. Trial sequentiality is a peculiar function of trial performance; as shown in (1B), not only higher trial sequentiality corresponds to higher performance but this increasing curve appears to have a distinctive form, with an inflection point in mid-performances.

If one reaches for the simplest explanations outside free association, those two seemingly unrelated facts appear to share some key features:

- In both phenomena, the extra mechanism at work with free association can be described as the avoidance of options in principle provided by free association – respectively (a) repetitious retrievals and (b) non-sequential retrievals.
- While the avoidance has different motives in the two cases and could be implemented in different brain areas, it may be assumed to operate in a de facto similar way, with each free association move being allowed or disallowed by a dedicated module depending on whether the retrieval meets a specific criterion (novelty or sequentiality).

These empirical observations lead to our working hypothesis, illustrated in Fig. 2. We will adopt the term “repetition bouncing” for the novelty-screening of free-association retrievals (i.e. the active avoidance of free association steps that would yield repetition, cf. Introduction) and will use the term “strategic bouncing” for the sequentiality-testing of free-association retrievals (i.e. the enforced avoidance of free association steps that would not obey a sequential ordering of recalls). We call it “strategic” as it is known that a bias toward sequentiality boosts performance (Zhang et al., 2021).

In the rest of the paper, we report on our detailed testing of the bouncing hypothesis from two complementary fronts – data mining and mathematical modeling. On the side of data mining, we exploit the large size of the dataset to tease out information not explicitly reported by individual subjects, with the aim of uncovering what has not been verbally outputted. On the side of modeling, we articulate the above intuitions as a streamlined theory that represents free recall without bouncing as a Markov process and bouncing as a non-markovian add-on; after fitting the corresponding parameters we will compare the predictions of the theory to observed features of the data.

### Model-free analysis of recall delays

If a mechanism exists by which the free association process is corrected, unwinded or rebooted, given that this prevents the outputting of some retrieved items, it will not leave direct traces within the record of recalled items. Yet, it may produced temporal delays in recall transitions where the process is activated. In both panels of Fig. 2 for example, the word “ear” will be recalled slower because first the word “toy” was retrieved and discarded. A natural approach is thus to hunt for traces of those delays in the recorded IRIs. And to do so, the first obligatory step is an a priori analysis of the main potential confounders.

Following convention (Kahana, 2012) we will call “output position” the ordered position of an item within the sequence of recalls reported during the recall stage. It can be a positive number if counted from the beginning of the recall process or a negative number if counted from the end (number of extant recalls to the end of the trial). For example, in the trial of Fig. 2A the word “sun” has output position = +2 or = −3. If trials with different performance are included, there is no fixed correspondence between positive and negative counting of outputs. It was shown in (Rohrer & Wixted, 1994) that the distribution of IRIs is characterized by negative output position. This is indeed apparent in the dataset under consideration, where dependence on the negative output position alone explains about 20% of the variance (Fig. 12). Note that the time interval from the onset of the recall stage to the first recall at output position = +1) is not an IRI but will be included for convenience among the IRIs and this does not qualitatively change our results.

Upon subgrouping all recall events by their negative output position, we compare events occurring before and after the item in a given serial position has been recalled (Fig. 3B). After recall of an item, its repetitious retrieval becomes possible. If such a retrieval comes to mind, it would be associated to a positive time delay. In other words, systematic *bouncing* against the item in a given serial position will be expected to make the post-recall IRI larger than the pre-recall IRI for every output positions (see Materials and Methods for details of the analysis). This turns out to be true for the last item in the list regardless of the output position. If the final item has already been recalled, all else being equal, a delay is found to occur in the recall process.

The fact that the final item appears to be a preferential bouncing target is a nontrivial manifestation of the recency effect (cf. Introduction). The delay associated to potential repetition of the final item can thus be thought of as a “silent recency effect“; if the attraction of the most recent memory from the list is the strongest, it can stay such after that memory has been recalled and subsequent recalls are slowed down through reversions to that item.

The absence of a corresponding silent primacy, on the other hand, is an unexpected finding. It may be related to the observation that recency typically appears earlier than primacy in the recall process (Fig. 10B) as the early recall of an item makes it the more likely to be bounced afterwards.

A similar analysis can be performed by comparing the conditions in which the retrieval transition with a given *serial-position lag* would pass or not the repetition screening. The mean IRI in these two conditions can again be compared, to check for a delay associated to the possible discard of such retrievals (technical aspects in Materials and Methods). As was mentioned in the Introduction, a widely ascertained property of reported recalls is that the most likely lag to occur is *L* = +1 (sequential recalls). If this type of transition remains the most probable one also when it can cause bouncing (i.e. when it would yield an item already recalled), we expect it to be associated to the largest delay.

The conjectures is confirmed by inspection of the data (Fig. 4). Thus, we find that another silent serial-position effect is the predominance of delays associated to the potential discarding of sequential retrievals (“silent contiguity effect”).

### A minimal model

To further test the bouncing hypothesis as applied to repetition avoidance, it is necessary to rely on forward models that describe it in a predictive fashion.

Models of free recall have covered a spectrum ranging from dual-store memory search models (Raaijmakers & Shiffrin, 1981; Raaijmakers, Shiffrin, et al., 1980) to powerful theories of temporal context dependence (Howard, Kahana, et al., 2002). It is not our ambition to provide a wide-ranging model of top-down control or in any way revise what is known from existing models (see Introduction); because our focus is entirely on the concept of bouncing, we rely on the most drastic simplification compatible with the general features of free recall.

The three assumptions we make to simplify the problem are the following (see Fig. 2A):

- in the absence of any bouncing, the reported recall process would be identical to the retrieval process and both would be a Markov chain in the space of the list items.
- in the presence of bouncing, the retrieval process can hit an undesired memory item and bounce back to the previously recalled item, and this step does not affect the sequence of reported items.
- every bounce entails a probability *q* of terminating the process, and we will refer to the parameter *q* as impatience.

Thus the only parameters are the Markov transition matrix and the impatience *q* (one additional parameter will be later introduced when we study strategic bouncing).

We have explored alternative assumptions for both the termination mechanism (static sink states, abrupt termination thresholds) and for the non-markovian add-on (one example is analyzed in detail in Supplementary Note 2), and they have proven less predictive than the model we are presenting.

The non-markovian contribution from bouncing against a recalled item is, in this framework, of a peculiarly simple type; namely, the transition probability to a given item becomes zero after that item has been recalled and stays zero up to the end of the recall process. This non-markovianity can be described as a progressive masking of the transition matrix.

The corresponding loglikelihood as well as its gradient (Supplementary Note 1) are easily written down and applied to fitting the dataset. This yields an impatience parameter *q* ≈ 0.1, and a transition matrix that encapsulates all known serial position effects including some degree of primacy (Fig. 13A). The number of bounces underlying every recall event can then be calculated by simulating the model with these parameters (for sample trials see Fig. 13B while the fitting and simulating protocols are documented in Materials and Methods).

To test the model we assume for simplicity that the IRI is proportional to the bouncing count. We first compare the mean of relevant variables characterizing a recall event. The dependence on negative output position, retrieved serial position, and the recall lag are all qualitatively recovered (Fig 5).

We then test for the silent serial-position effects along the same lines followed for real data (Figs. 3-4), with the only difference that instead of measuring a mean temporal delay between two conditions, we now measure a difference in the mean number of bounces occurring within a recall event. We report that all main phenomenological features of silent serial-position effects, discussed above, are found in our pseudodata, both as concerns silent recency (Fig. 6) and silent contiguity (Fig. 7).

### Model-free analysis of sequential slowdown

We turn then to the distinctive shape of the sequentiality/performance curve (Fig. 1B). We hypothesized that it might emerge if a partial discarding of free-association retrievals allows the implementation, in some trials, of a more markedly sequential recall strategy associated to higher performances – yielding the upturn of the curve for high-performing trials. (This interpretation would also justify the related fact that the output position of the final item becomes bimodally distributed for highly performing trials, Fig. 14). If that were the case, it should also leave traces in the statistics of the IRIs. Namely we would expect that, at least among high performers, recalls performed by strategizing would be slower than those performed without.

**Figure 1.**
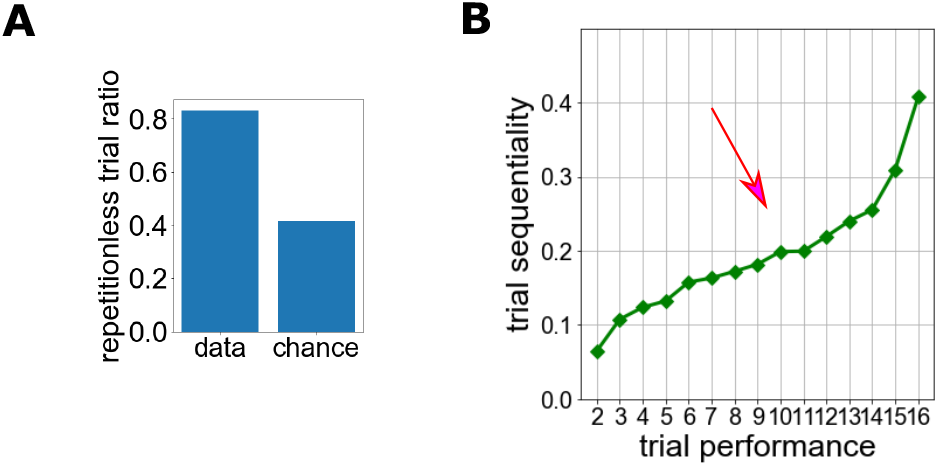
Two features of free recall. **(A)** Paucity of observed output repetitions as compared to chance level (for details of chance level estimation see Fig. 11). **(B)** Fraction of sequential recalls in a trial as a function of trial performance (i.e. number of recalled items). Mean curve is shown, while the standard error of the mean is too small to plot. Red arrow highlights approximate location of an inflection point, where the slope starts to increase.

**Figure 2.**
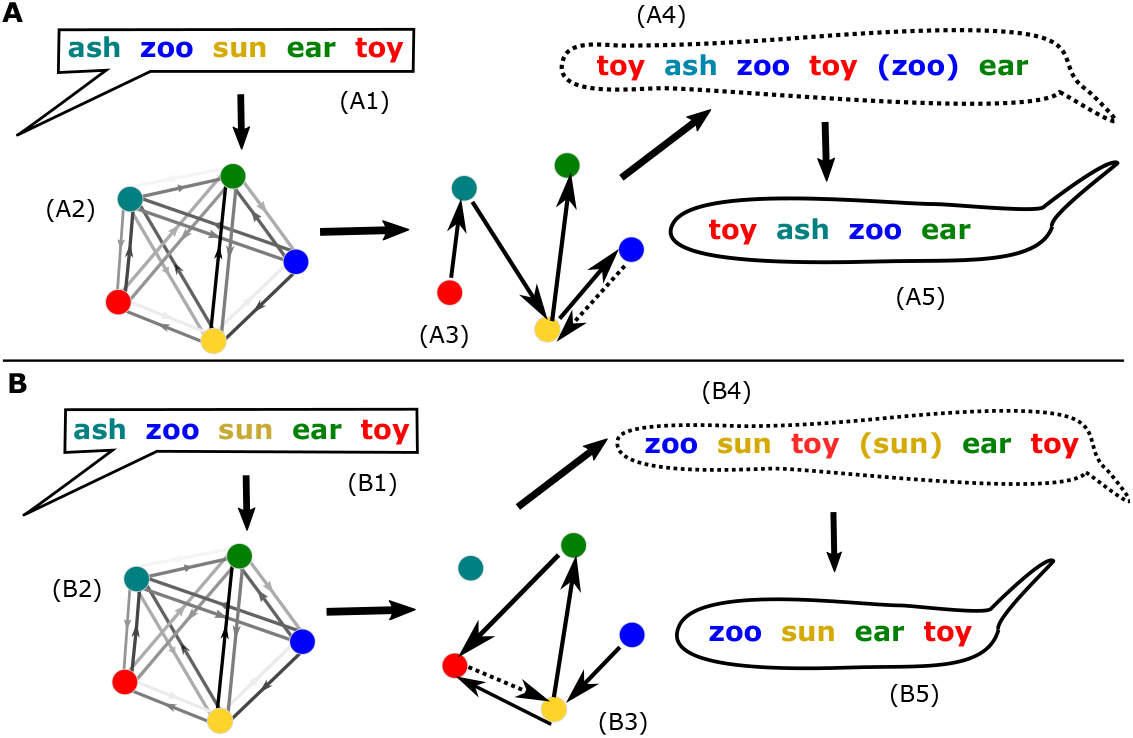
Schematic depiction of bouncing. Intervention over the free association process by “bouncing” simultaneously explains paucity of observed repetitions and higher sequentiality of high-performance trials. Here, a five-item list is presented in two separate trials leading respectively to an instance of *repetition bouncing* (**A**) or one of *strategic bouncing* (**B**). During the presentation stage of free recall (A1,B1) a subject is presented visually or orally with a list of *L* items (here *L* = 5 words). Memories of exposure to each of the items are stored in such a way that mental transitions among memories have unequal probabilities represented in (A2,B2) by the shades of gray of the arrows. During the recall stage (A3), the subject instantiates a mental trajectory through those memories(A3,B3). Nothing prevents the sequence of retrieved memories from containing an unlimited number of repetitions (as for example in A3,A4), but the output is devoid of repetitions (A5), posing the problem whether some retrievals are discarded. In addition, a tendency toward sequentiality in high-performing trials is also observed (B5), posing the problem whether non-sequential transitions have been actively avoided (as when stepping back from “toy” to “sun” in B3,B4). Bouncing transitions are displayed as dotted arrows.

**Figure 3.**
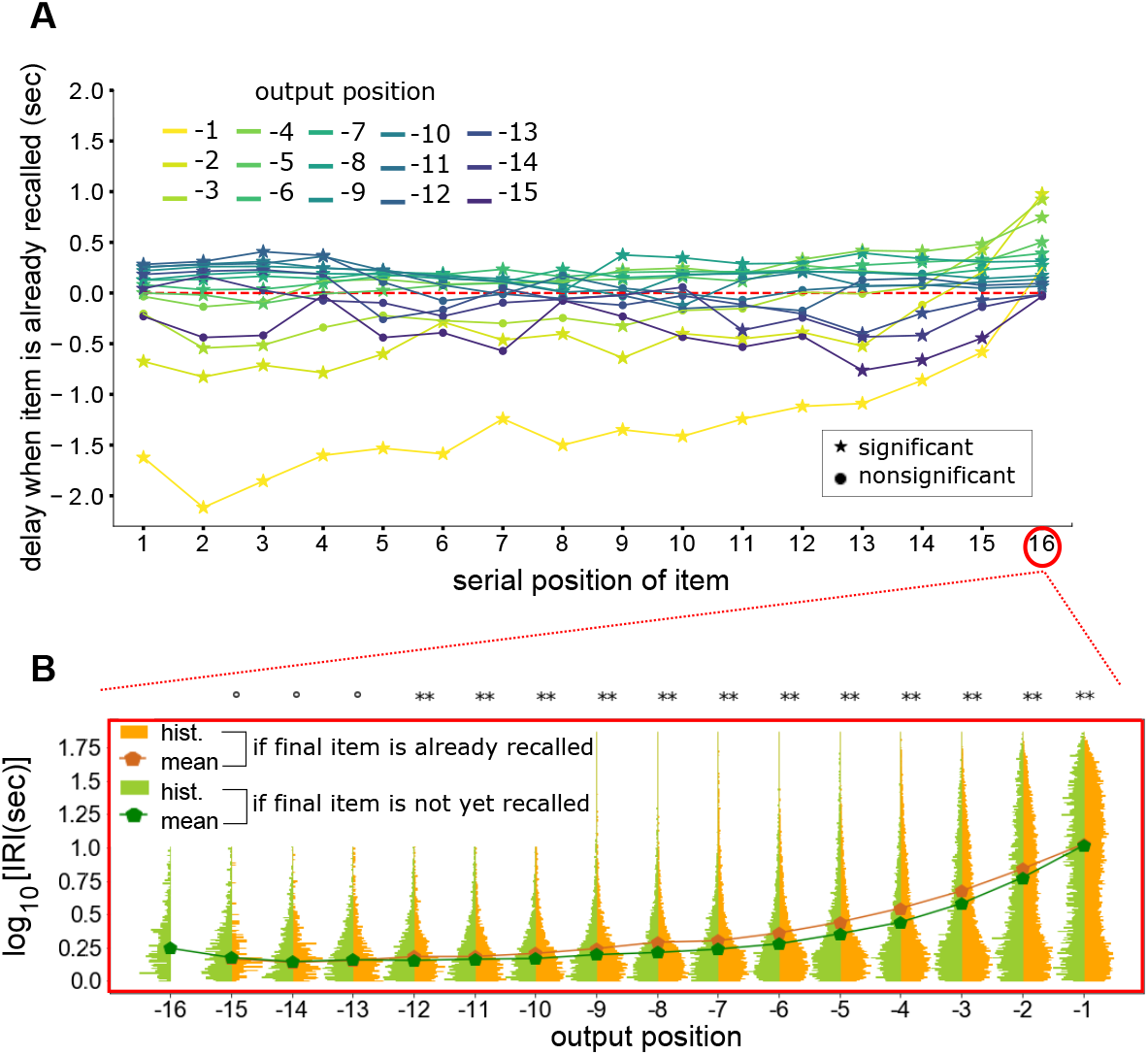
Empirical evidence for a higher-order “silent recency effect” in free recall. **(A)** Difference between mean IRI when subsampling on the basis of whether the item with a given serial position on the list has already been recalled or not. IRI were histogrammed excluding recalls of the item with the given serial position also when it is not yet recalled. The serial position concerned is shown on the x-axis, and plot markers encode significance as defined by a Mann-Whitney u-test p-value lower than 10^−3^ (star = significant, circle = nonsignificant). **(B)** Histogram of the IRIs for recalls of items other than the final one, plotted separately for each output position and for the two conditions were the final item has already been recalled or not. The scale of the IRIs is logarithmic. P-value from the Mann-Whitney u-test is encoded as follows: circle, *p* > *p*\* = 10^−3^; one star, *p*\* = 10^−3^ > *p* > 10^−4^; two stars, 10^−4^ > *p*. The previously occurred recall of the final item is statistically associated to a delay in the recall of other items.

**Figure 4.**
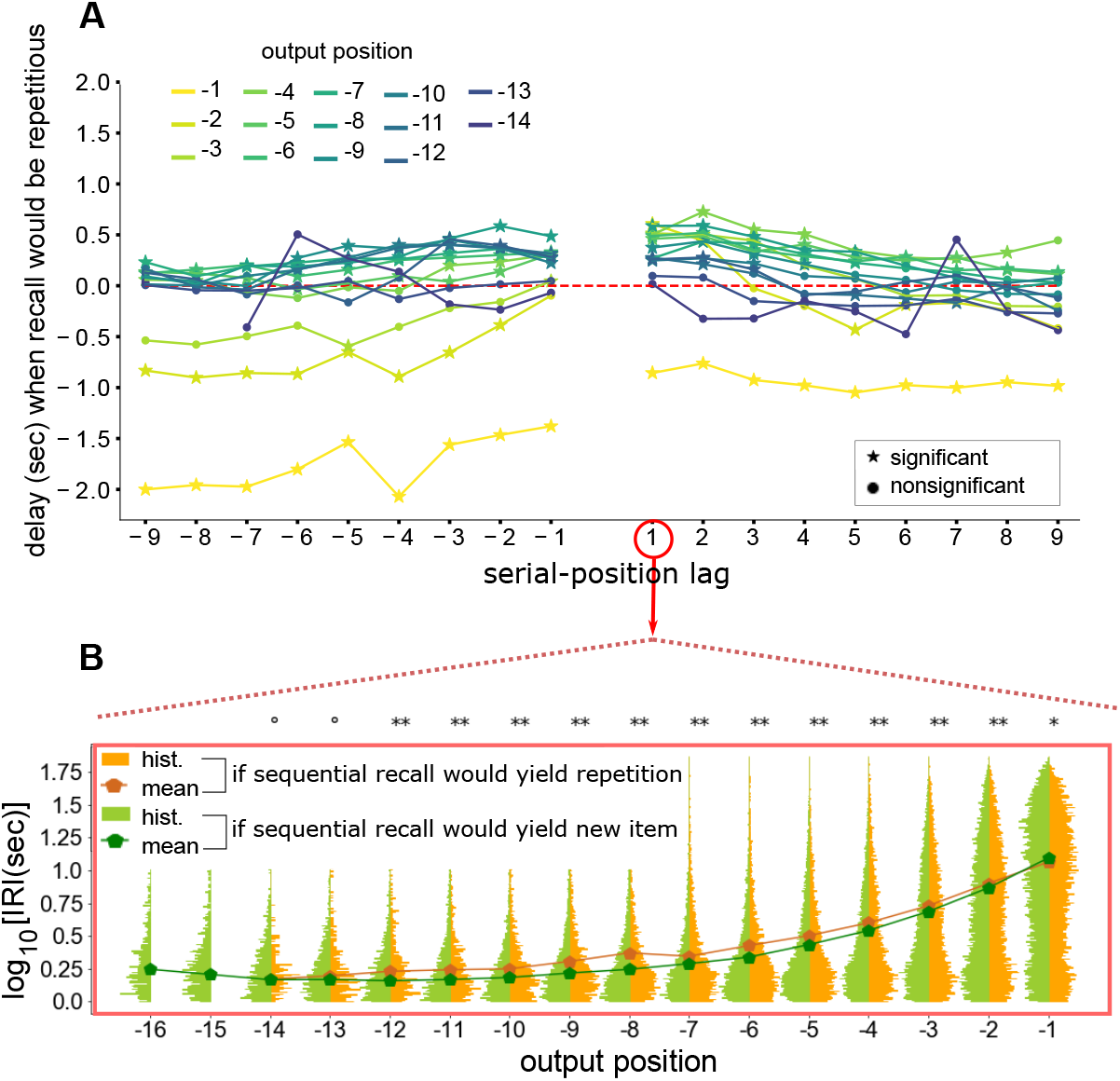
Empirical evidence for a “silent contiguity effect”. **(A)**: Difference between mean IRI when subsampling on the basis of whether the item situated at a certain fixed lag from the last recall has already been recalled or not. The lag position concerned is shown on the x-axis, and plot markers encode significance as defined by a Mann-Whitney u-test p-value lower than 10^−3^ (star = significant, circle = nonsignificant). **(B)** Histogram of the IRIs for recalls of items other than the sequential one (i.e., the one that was presented right after the latest recalled item) shown separately for each output position and for the two conditions were the sequential item has already been recalled or not. The scale of the IRIs is logarithmic. P-value from the Mann-Whitney u-test is encoded as follows: circle, *p > p*\* = 10^−3^; one star, *p*\* > *p* > 10^−4^; two stars, 10^−4^ > *p*. The previously occurred recall of the sequential item is statistically associated to a delay in the recall of other items.

**Figure 5.**
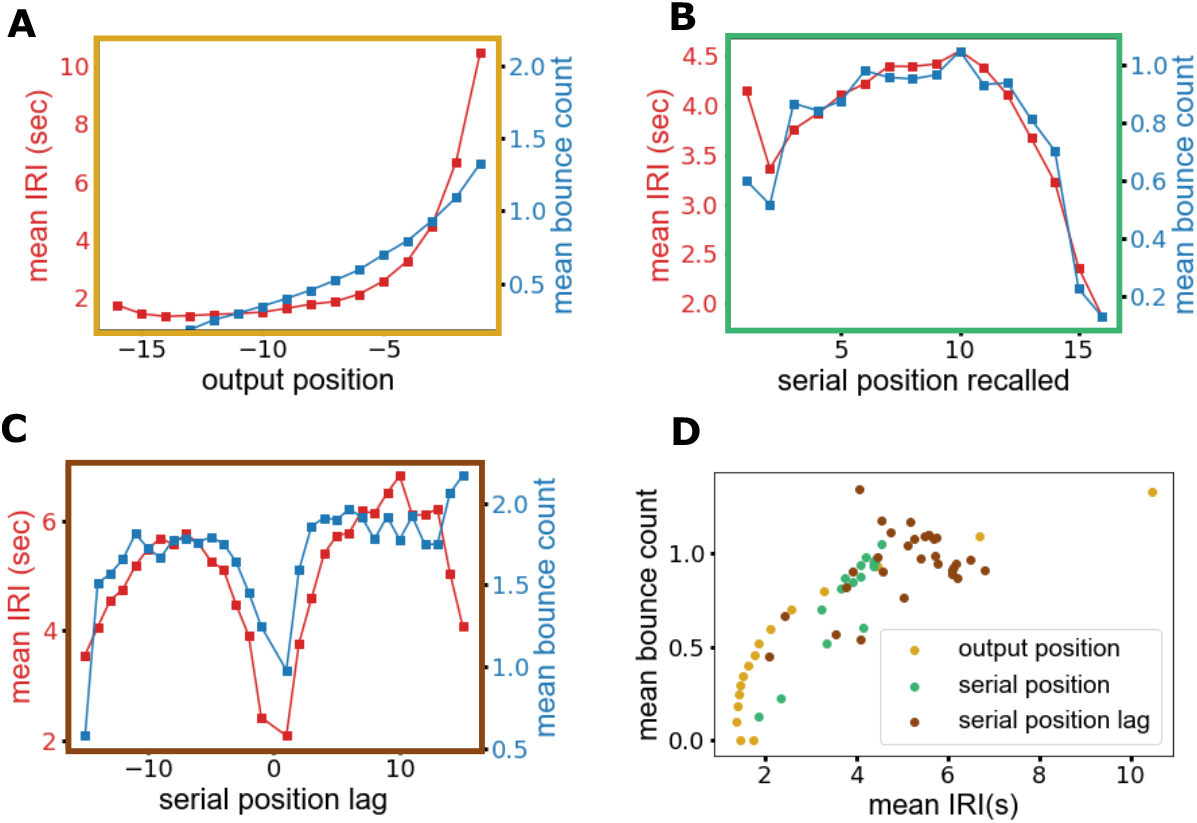
Comparison between mean IRIs and the mean number of unreported retrievals of the model. Result of averaging is shown for fixed values of the recall event’s output position **(A)**, the serial position of the recalled item **(B)**, and the serial position lag from the preceding recall **(C)**. In **(D)**, values shown in the previous panels have been scatter-plotted together.

**Figure 6.**
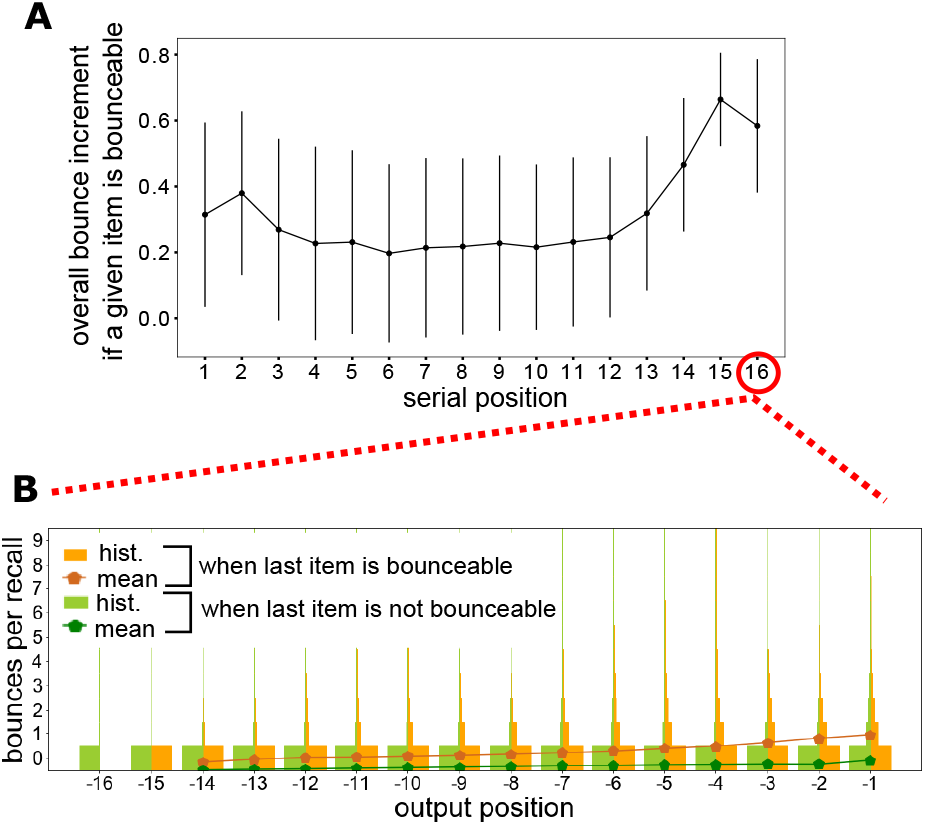
Silent recency effect emerging from the bouncing model. **(A)** Selective variations in the mean bouncing counts estimated from model simulations. The mean number of bounces within a recall event has been averaged in the two subsamples of recalls where the list item in a given serial position is or is not bounceable, and the difference between the two means is plotted as a function of the serial position. Values on the x-axis refer to the item by the bounceability of which subsamples are discriminated. **(B)** Bouncing-count histograms displayed side by side refer to recall events where the final item in the list is or is not bounceable. Corresponding mean curves show the mean bouncing increment after recall of the final item.

**Figure 7.**
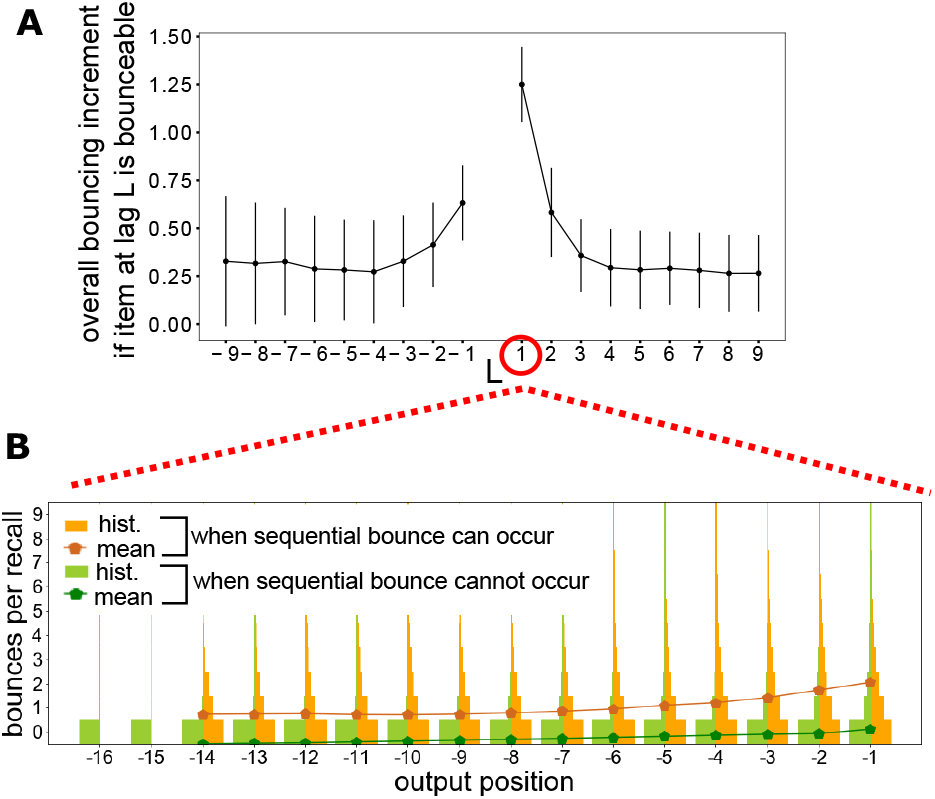
Silent contiguity effect emerging from bouncing model. **(A)** Selective variations in the mean bouncing count per recall estimated from model simulations. The mean number of bounces within a recall event has been averaged in the two subsamples of recall events where a given lag does or does not lead to a bounceable item, and the difference between the two means is plotted as a function of the given lag. **(B)** Bouncing-count histograms displayed side by side refer to recall events where the sequential candidate for recall is or is not bounceable. Corresponding mean curves show the mean bouncing increment when the sequential item was already recalled.

To differentiate between strategic and non-strategic trials we can use the fact that the former have by construction a higher trial sequentiality. An obvious approach is to separate each given sample of trials in the submedian and supramedian subsamples according to its distribution of trial sequentialities, and test whether recalls are faster in the submedian subsample.

We take into account three confounders. First, the dependence of the IRI on the negative output position can again play the role of a strong confounder. Second, since sequential strategies are used to achieve higher performances (see Introduction), the effect we are seeking may only be present in the more highly performing trials. We must thus proceed by considering separately event sets defined both by a given output position and by their occurring in trials with given performance. The third confounder is the difference of IRIs for sequential and non-sequential recalls within each trial. Indeed, sequential recalls are mostly faster than non-sequential ones across different performance levels and output positions (Fig. 15A).

For every condition defined by trial performance and output position, we consider thus the distribution of trial sequentiality among trials with that performance that contain a sequential recall in that output position (Fig. 8A). We then split the sample by the median value of trial sequentiality, and compare the IRIs of recall events in the two subsamples (Fig. 8B). We only compare sequential recall events because highly sequential trials contain few non-sequential recalls. Thresholding out all sample pairs that do not pass a significance test, we calculate the sign of the delay associated to higher trial sequentiality. (Results are qualitatively unvaried whether we partition using a median computed over the sequentialities of all trials in a performance group or computed separately over those associated to sequential events at each given output position.)

**Figure 8.**
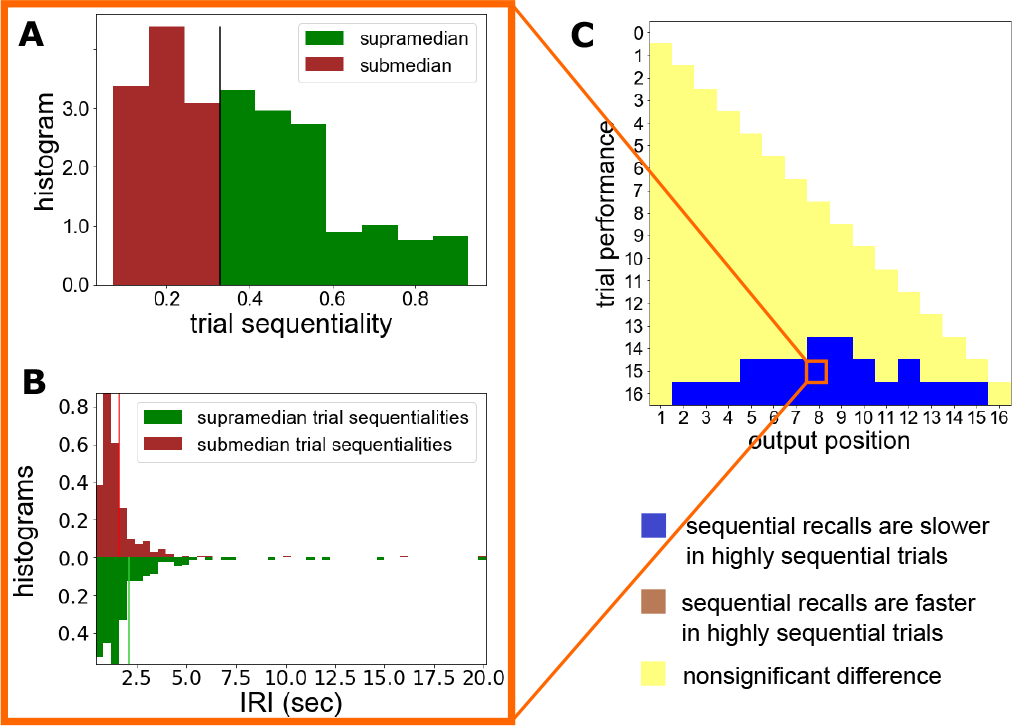
“Sequential slowdown” in free-recall data. For each performance level, we list all trials that perform a sequential transition at a given output position and study how its speed changes in highly sequential trials. The sequentiality value associated to each trial is simply the fraction of sequential recalls in the trial. **(A)** The histogram of this quantity for a particular subsample is further subdivided for the sake of the analysis into the set of trials above and below its median. **(B)** The corresponding distribution of IRIs in the two subsamples, with vertical lines indicating the respective mean values. **(C)** Sign of the difference in the mean values whenever it is significant by a Mann-Whitney test with threshold *p*\* = 10^−3^. Sequentially biased trials are characterized by a slowdown of their sequential recalls, specifically in high-performing trials. Squares on the grid are colored as follows: *white* if data are not available from both subgroups (as is the case here because output position *K* does not exist for trials with performance < *K*); *yellow* if the Mann-Whitney p-value between the two corresponding subsamples is larger than *p*\* = 10^−3^; *blue* if the *p* > *p*\* and the difference is positive, i.e. the supramedian trials perform significantly slower than sequential recalls; the opposite occurrence would be submedian trials performing significantly slower on sequential recalls but, strikingly, this is never observed.

We thus find that (see Fig. 8C):

- In all cases where our significance requirement is met, the delay is positive, meaning that sequential recalls occurred in sequentially biased trials have taken longer. In the following we refer to this phenomenon as *sequential slowdown*.
- Strikingly, the phenomenon is concentrated in the high-performance regime. This is in fact just where it is to be found if the sequential slowdown is due to the implementation of recall strategies.

We will focus on the hypothesis that the slow sequential recalls are due to a subselection of the retrievals (for the problem with alternative explanations, see Discussion) or, equivalently, these delays are punctuated by the active rejection of what non-sequential retrievals are provided by free association, which is what we termed strategic bouncing.

### Sequential slowdown in the bouncing model

With model parameters coming from the above-mentioned fit, we only add into the model a finite probability *s* (“strategicity”) of bouncing back from a retrieved item if it is non-sequential. We are not necessarily assuming a common mechanism for strategic bouncing and repetition bouncing (the fact that strategic bouncing happens only in a minority of trials may point to the opposite) but only that the basic retrieval rejection step can also be described as a bounce. A priority needs to be established between the two types of bounce; we assume that subjects who are actively implementing a sequential strategy check first for sequentiality and then for repetitiousness and do not associate any impatience increment to strategic bounces. This proves to be the sensible choice from the requirement that the performance should increase as a function of strategicity (Fig. 9A) despite the recency effect (Fig. 10).

**Figure 9.**
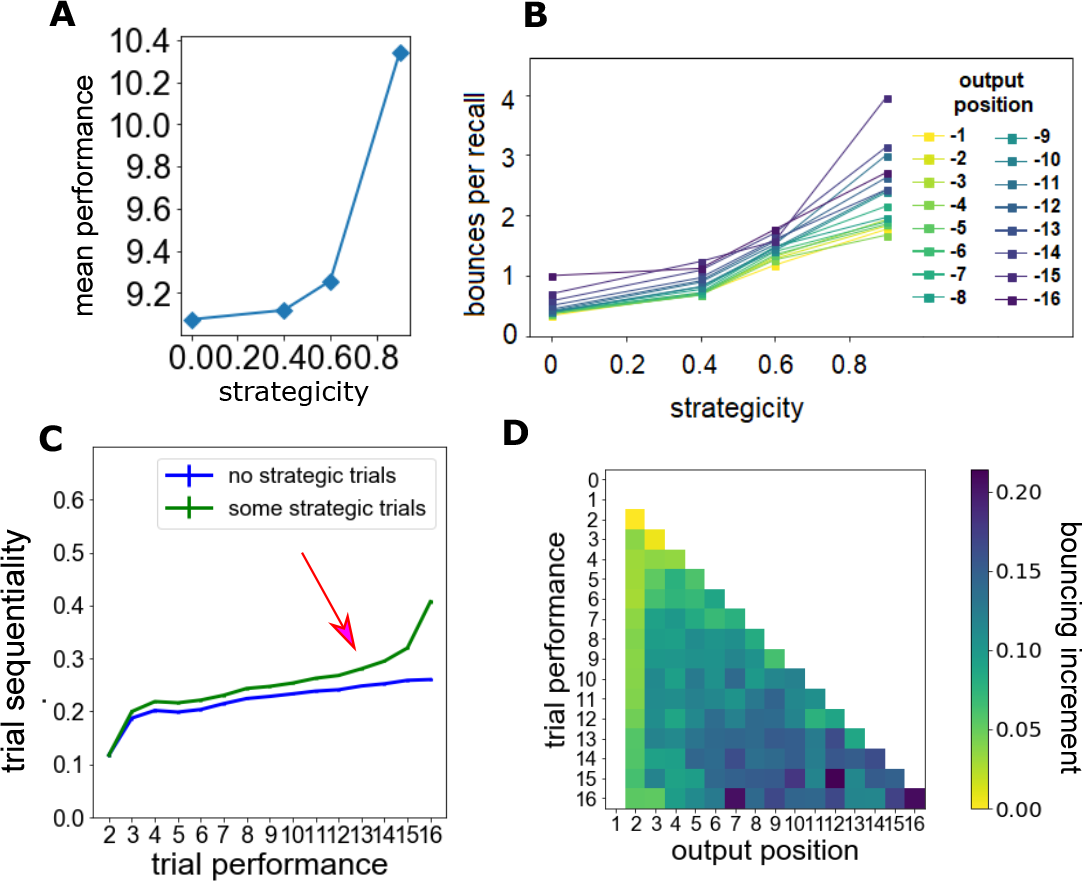
Simulations of the bouncing model with strategizing. We simulated the model by adding in a given probability of strategic bouncing (i.e. the probability of rejecting a non-sequential retrieval), which we termed “strategicity”. **(A)** Increasing strategicity also increases performance, coherently with the theory. **(B)** Increasing strategicity slows up the process, which agrees with the slowdown of sequential recall in highly sequential trials seen in the data. **(C)** The introduction of strategic bouncing leads to the emergence of the inflection point (red arrow). The features of real data that were shown in Fig. 1B are recovered here by merging a dominant population without strategicity, *s* = 0 (5 × 10^5^ trials corresponding to the blue curve) with a smaller one having strategicity, *s* = 0.5 (5 × 10^3^ trials). **(D)** Here, for each joint condition of performance and output position, sequential recalls are sampled above and below the median value of trial sequentiality. The mean difference in bouncing count proves to be always positive and increases in the area where sequential slowdown was detected in the data (compare Fig. 8D).

**Figure 10.**
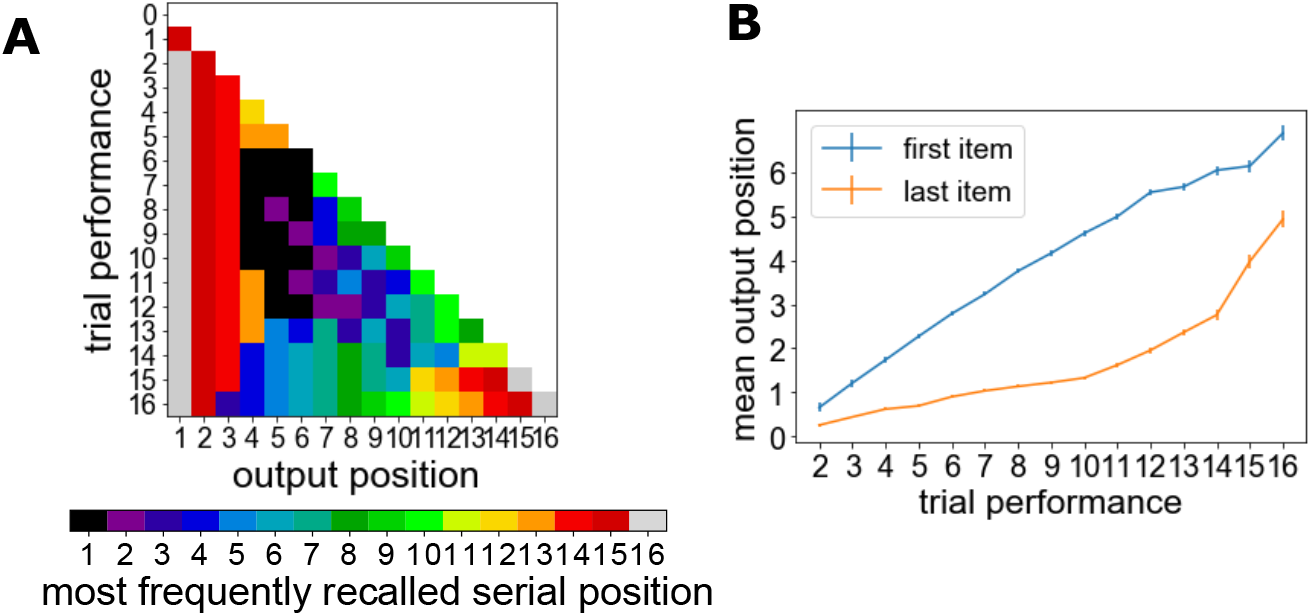
Overview of recall by serial position of recalled items. **(A)** Most commonly recalled serial item at any given output position in trials with given performance. Notice in particular the bold recency effect, the subsequent primacy cluster, and the anomalous termination of the high performance trials. **(B)** Mean output position of first and last serial item, highlighting that primacy effects intervene after recency effects on average (error bars show the standard error of the mean).

The analysis of sequential slowdown in data suggests that this increase in performance should come at the price of a slowdown in individual recalls, measured by the number of bounces. We test this statement on simulations of the model and find that indeed it holds true for simulated recalls with any possible value of their negative output position (Fig. 9B).

We then move on to replicating the results of data analysis for the dependence of trial sequentiality on performance that was presented at the outset (Fig. 1B). Rather than aiming at numerical accuracy we seek to understand the shape of the curve, namely its inflexion in mid-performances, followed by a drastic rise for larger performances. First we note that repetition bounces alone don’t explain this feature (Fig. 9B). We then simulate a mixture of two populations, a dominant one without strategicity, *s* = 0 (5 × 10^5^ trials), and a smaller one with strategicity, *s* = 0.5 (5 × 10^3^ trials). When plotting the sequentiality of simulated trials versus their performance, we find that even such small percentage of strategic trials is sufficient to inflect the curve (Fig. 9B).

Finally, we agnostically interrogate the mixed-set pseudodata model for the signatures of strategic slowdown we encountered in Fig. 8C. We subsample recall events by the output position and trial performance characterizing them; for each joint condition, we use the sequentiality values of the relevant trials to subdivide the sample in a supramedian and submedian group and compute the bouncing increment, i.e. the difference between their mean bouncing counts.

This difference proves to be always positive (Fig. 9D), signifying a delay in time as found in real data. In Fig. 8C we also showed that the time delay found in real data grows into significance in the high performance region for sufficiently large output positions. The model will be in agreement with real data if, whenever the data show a statistically significant difference in IRIs, the corresponding subsample of the model-generated pseudodata features relatively high values of the bouncing increment. This prediction is finally confirmed by the result of Fig. 9D further supporting the notion that sequential slowdown is due to strategic bouncing.

## Discussion

While free recall experiments are an essential behavioral probe in the study of human memory, much research has revolved around standard serial-position related effects (see Introduction). In this work, we undertook the rigorous mining of a large dataset to uncover a number of features:

- the number of repetitions observed in these experiments is vastly smaller than what a basic expectation given by chance Markovian transitions;
- recalls are delayed if the final item has already been recalled (“silent recency effect”);
- recalls are delayed if sequential recall would yield a repetition (“silent contiguity effect”);
- the overall abundance of sequential recalls in the trials has a distinctively inflected shape when regarded as a function of trial performance;
- a slowdown in certain recall events is associated to the overall prevalence of sequential recalls in the trial (“sequential slowdown”).

In addition to reporting on the above, we proceeded to suggest a unifying explanation coherent with the current body of knowledge on free recall (see Introduction), and based on the notion that free-association retrievals are sometimes “bounced”, i.e. discarded. In a majority of cases this happens because they are repetitious (repetition bouncing) and in a minority because they are not sequential (strategic bouncing).

To test the validity of our explanation we used a minimal model with trajectories driven by heterogeneous transition probabilities and reset by bouncing. In terms of its complexity, this model of free association is situated halfway between the extremes of the pure-death model discussed in the Introduction (incapable of modeling contiguity effects, see McGill (1963)) and sophisticated models of free-associating neural networks such as Romani, Pinkoviezky, Rubin, and Tsodyks (2013).

In the model, bouncing occurs systematically for repetitious retrievals and adds to the “impatience” that, probabilistically, leads to termination of a recall trial; bouncing of nonsequential transitions occurs in strategic trials with a fixed probability (“strategicity”). Fitting these parameters on data and using the number of bounces as a proxy for the inter-recall time intervals (IRIs), the model was shown to qualitatively account for all the above features of free recall.

Our work underscores the importance of unreported retrievals in determining behavioral observables of memory experiments. It is easy to see indeed that alternative explanations would run into several problems.

For example, one might hypothesize that the strategy at play in certain high-performing trials consists merely in waiting longer before surrendering even if no new memory is being retrieved. This hypothesis predicts a slowdown preferentially happening at later output positions, where surrendering is most prominent. However, such explanation can be ruled out because the sequential slowdown is observed at a wide range of output positions (Fig. 8C).

Neither our analysis nor our models have relied on specific assumptions on the source of the post-retrieval control, although active blocking of internally recalled items may be conjectured to emerge from higher-level processes that operate top-down over free association (see Introduction). However, our results can be taken as a starting point to tackling that question.

Our statistical inquiry on the IRIs dove to a considerable level of detail but still relied exclusively on mean values of the IRI over different subsamples. The full density profiles of subsampled IRI distributions contain a wealth of unexplored information that deserves to be the subject of a separate study.

A key advantage of the minimal model we employed is how easily it accounts simultaneously for repetition bouncing and strategic bouncing. One disadvantage is that it cannot account for the gradual release from repetition avoidance observed in data from both free and serial recall (Henson, 1998; Laming, 2010; Rundus, 1971). This could be accounted for in future work using what is known about its dependence on cognitive rather than physical time (Duncan & Lewandowsky, 2005).

Measurements performed in free recall are not limited to behavioral observables and numerous studies have been performed using noninvasive neuroimaging techniques such as EEG, MEG, PET, fMRI (Brassen, Weber-Fahr, Sommer, Lehmbeck, & Braus, 2006; Dickerson et al., 2007; Norman et al., 2017; Savage et al., 2001; Staresina & Davachi, 2006) as well as invasive methods in epilepsy patients (Gelbard-Sagiv, Mukamel, Harel, Malach, & Fried, 2008; Solomon, Lega, Sperling, & Kahana, 2019). A parallel study of unreported retrievals could be attempted through the decoding of suitable neurophysiological data. The timing and magnitude of post-retrieval effects such as those already observed in ECoG of prefrontal cortex (Norman et al., 2017) could be compared to the timing and quantity of the bounces inferred by an analysis of behavioral data.

The issue whether bouncing is consciously perceived by the participants is more delicate, and it could be partly addressed by comparison with Externalized Free Recall (Hogan, 1975; Kahana et al., 2005; Roediger & Payne, 1985), with the caution of providing separate controls as for the accuracy and consistency of self-reporting.

In this work we focused on the two types of bounces – repetition bouncing and strategic bouncing to maintain sequentiality – merely because these two were compellingly suggested by Fig. 1. In different datasets on human recall the criteria that drive post-selection will accordingly be different but are bound to leave detectable traces in the IRI statistics, specifically with tasks where some recalls are to be expressly excluded. For example in Roediger III and Tulving (1979) subjects are instructed to recall items not belonging to a specified categories or items not beginning with certain letters. While other types of bouncing might take place, their detection can be attempted by the very means we have introduced.

The features uncovered by our model-free analysis of data should be recoverable with little adjustment from a variety of models such as the generally applicable framework of CMR (Lohnas et al., 2015; Polyn et al., 2009) or the stochastic model of Laming (2009). Mechanistic models based on neural networks with overlapping attractors (Romani et al., 2013) have also recently emerged as capable of remarkably accurate experimental predictions on free recall (Katkov, Romani, & Tsodyks, 2017; Naim, Katkov, Recanatesi, & Tsodyks, 2019) and involve already some degree of post-retrieval editing as part of their termination mechanism (based on the subject’s ability to detect certain retrieval loops and stop outputting). What remains to be understood is how the top-down post-retrieval action we described here as bouncing is effectively implemented by a neuronal network. A natural candidate would be inhibitory control (response inhibition), and testing its implementation on a neuronal network will be a further step toward unraveling the biological substrate for post-retrieval operations.

In conclusion, our data mining approach has shed light on the range and robustness of unreported cognitive processes behind memory recall. We showed moreover that a single minimal model could account for multiple types of post-retrieval mechanisms. Overall, our findings contribute to placing on a firmer foundation the evidence that post-retrieval mechanisms are a potentially ubiquitous ingredients of human memory.

## Materials and Methods

### Data

We base our analysis on a publicly available dataset, collected as part the Penn Electrophysiology of Encoding and Retrieval Study at the University of Pennsylvania (for a full account see Healey et al. (2014)). Participants consented according the University of Pennsylvania’s IRB protocol and were compensated for their participation.

Each word was drawn from a pool of 1,638 words. Lists were constructed such that varying degrees of semantic relatedness occurred at both adjacent and distant serial positions in a way that randomized semantic effects.

For each list, there was a delay of 1.5 seconds before the first word appeared on the screen. Each item was on the screen for 3 seconds, followed by a variable time gap lasting between 8 and 12 seconds (uniform distribution). The recall stage began after 12–14 seconds from presentation of the last item.

Participant were given 75 seconds to attempt to recall aloud any of the presented items. Because of the limited amount of time, quantities pertaining to the first negative output position are indeed somewhat affected by the final rush to recall.

In our analysis all trials (27198 in total) were used. In the minority of trials containing some repetition (Fig. 11A), repetitions were ignored for the main analyses. Intrusions from outside the presented lists were all removed at the outset.

**Figure 11.**
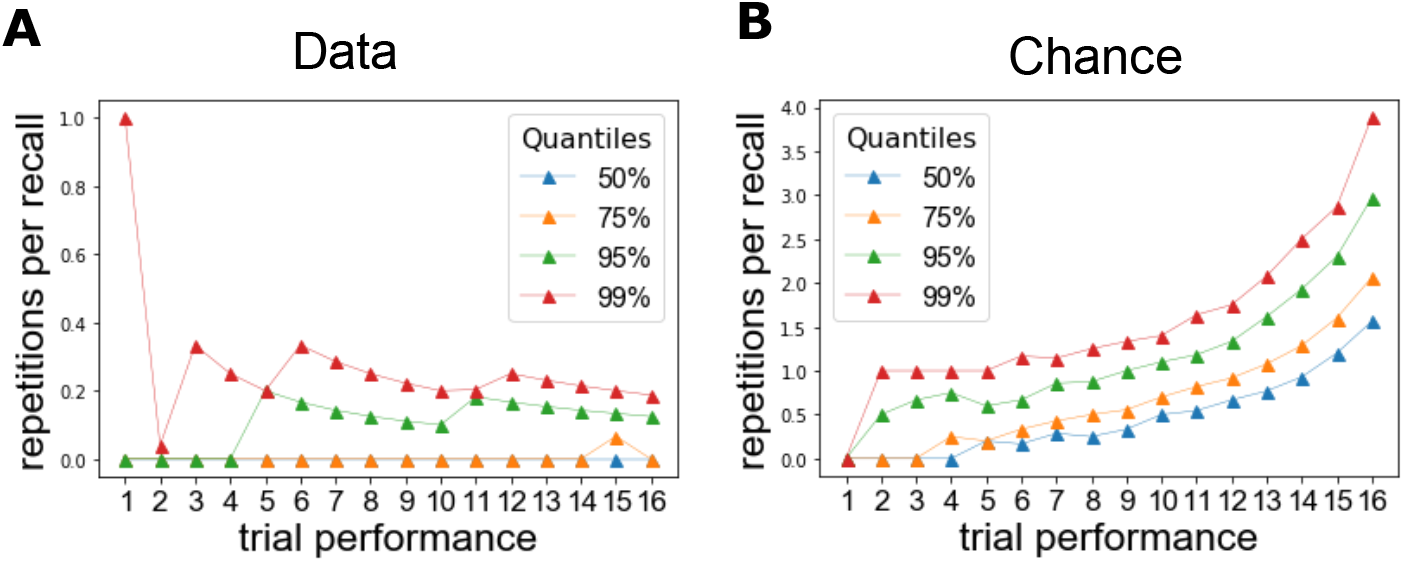
Comparison to chance level of the amount of repetitions in the dataset. **(A)** Statistics of repetitions per trial. If we subgroup trials by their performance, we find that the number of repeated recalls increases as a function of performance but even for large performance over three quarters of trials contain no repetition, and the others little more than one each. **(B)** To provide a quantitative baseline, we simulated a purely associative model, which consists of Markovian transition in between serial positions. Notice that the model is complex enough to accommodate heterogeneous transition probabilities. Since this model does not incorporate bouncings, termination cannot be induced through the “impatience” mechanism discussed in the main text. Thus, we counted termination as transition to a sink state so that the transition matrix driving the process is of size (*N* + 1) × (*N* + 1). We extracted this transition matrix from a fit of the dataset and ran it starting from an initial condition (distribution of the first recall over serial positions) directly extracted from the dataset. The number of resulting repetitions is vastly larger than in the data.

### Statistical Analyses

The set of potentially unreported retrievals was characterized by either the serial position of the retrieved item (Fig. 3) or the lag from the latest recall to the item (Fig. 4). In both cases, to make the analysis of IRIs rigorous we took the following precautions:

- We broke down the sampling of recall events by negative output position to discount the strong output position dependence highlighted in Fig. 12.
- We confronted pre- and post-recall conditions by comparing only the IRIs for the recall of items outside the putatively unreported set. Reported recalls of an item from that set can appear in the pre-recall sample but not in the post-recall sample, so we excluded them from both.
- In the case of unreported retrievals with a fixed lag, we count out all conditions where the lag would not yield an item within the list (due to the finite list length), as that makes recall unavailable rather than bounceable.

**Figure 12.**
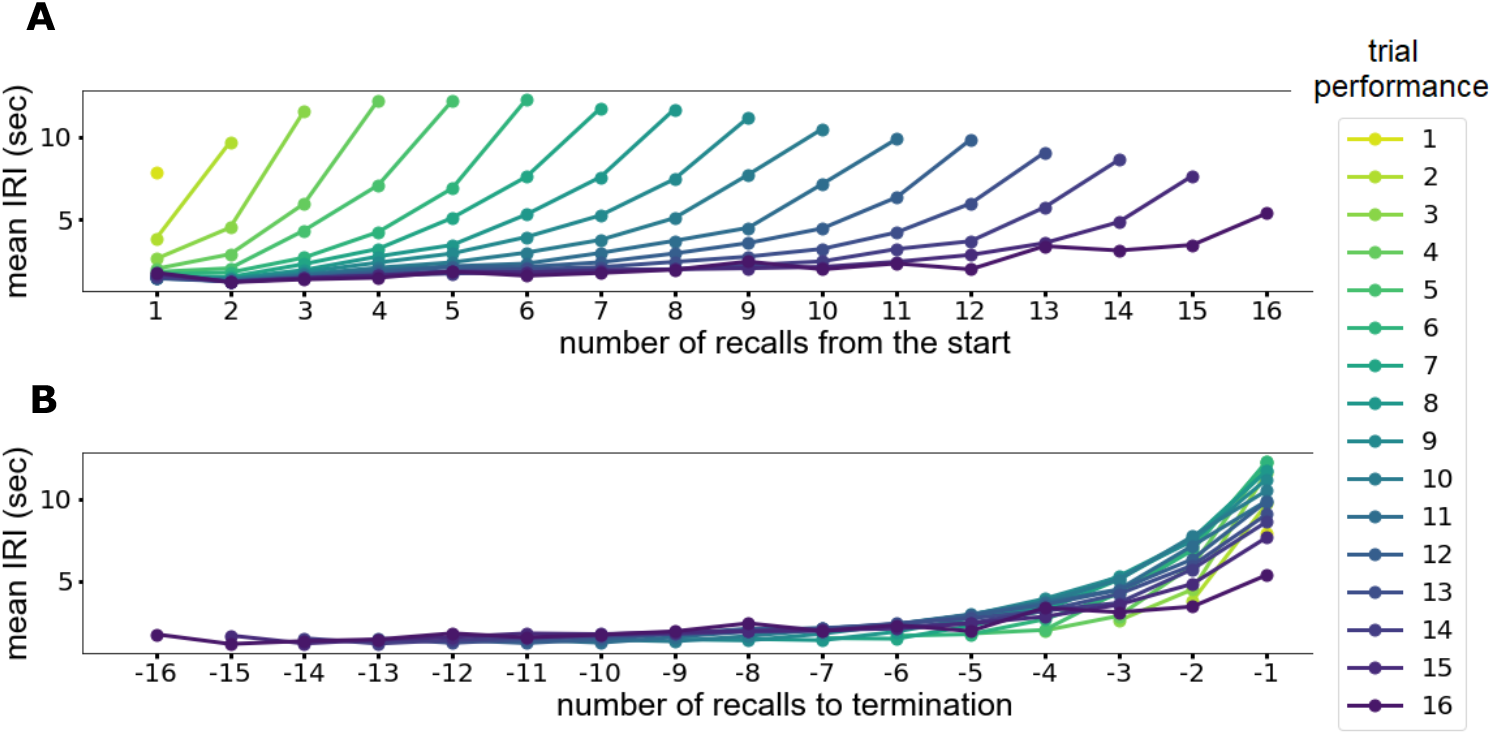
Terminal Alignment of mean IRIs. **(A)** Mean IRIs computed over trials with fixed performance. Aligning the curves by the first recall of the trials does not reduce observed variance. **(B)** Alignment at the end, on the other hand, makes the mean IRIs collapse (and explains 20% of variance in the IRIs).

In all the subsample comparison of the IRIs, we systematically filtered out information by testing for significance the difference between the IRIs in each pair of subsamples. Since the distribution of IRIs in any given subsample is highly non-gaussian (see e.g. Fig. 3A), rather than using a t-test we adopted a Mann-Whitney u-test (Hettmansperger & McKean, 2010).

For example, Fig. 15A should be strictly understood as stating that if we pool trials regardless of their sequentiality, the statement that sequential recalls are faster withstands a significance test with few exceptions on the triangular grid spanned by performance and output position values.

The p-value threshold we chose at the outset was *p*\* = 10^−3^ and has been uniformly applied to all the analyses.

### Model Fitting

Let 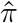 be the naked transition matrix (the zero-diagonal Markov transition matrix of the bounce-free model) with matrix elements *π*(*y*|*x*) determining the transition probability from item *x* to *y* and, for any set *S* of serial positions, let *π*(*S*|*x*) = ∑_*y*∈*S*_ *π*(*y*|*x*).

Further naming *N* the length of the presented lists, *T* the number of trials in the sample, *m_α_* the performance of trial *α*, *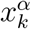* the serial position of the *k*-th recall in the trial, *q* the impatience parameter, and using the notation where *x_i:j_* is the set of serial position recalled in between output positions *i* and *j*, both included, the normalized loglikelihood of the bouncing model is

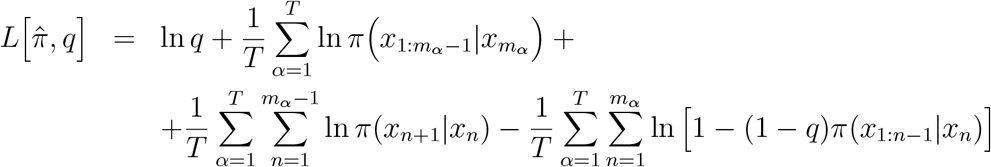

For the derivation, Supplementary Note 1.

We write down the gradient (Supplementary Note 1) and search for the minimum from a random initial condition using the L-BFGS-B algorithm (Byrd, Lu, Nocedal, & Zhu, 1995). This converges in < 10^2^ iterations to the matrix shown in Fig. 13A and to the optimal impatience *q* = 0.1.

**Figure 13.**
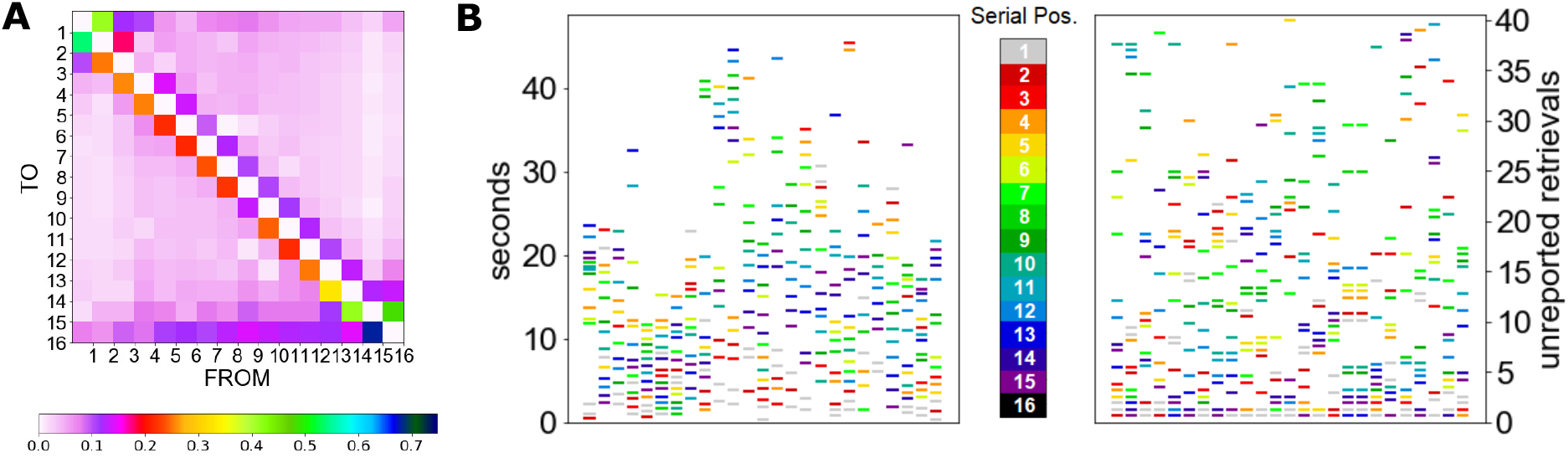
Illustration of typical features of the bouncing model. **(A)** The optimal transition matrix obtained from fitting the data with the optimal impatience *q* = 0.1. **(B)** Example trials from the data (Left) and from simulations of the bouncing model (Right). Color encodes serial position of recalled items. The y-coordinate in the data panel is the timing of the recall event. The y-coordinate in the model panel is the number of unreported retrievals observed up to a given recall in simulations of the model, which has been jittered to prevent overlaps for visual aid.

**Figure 14.**
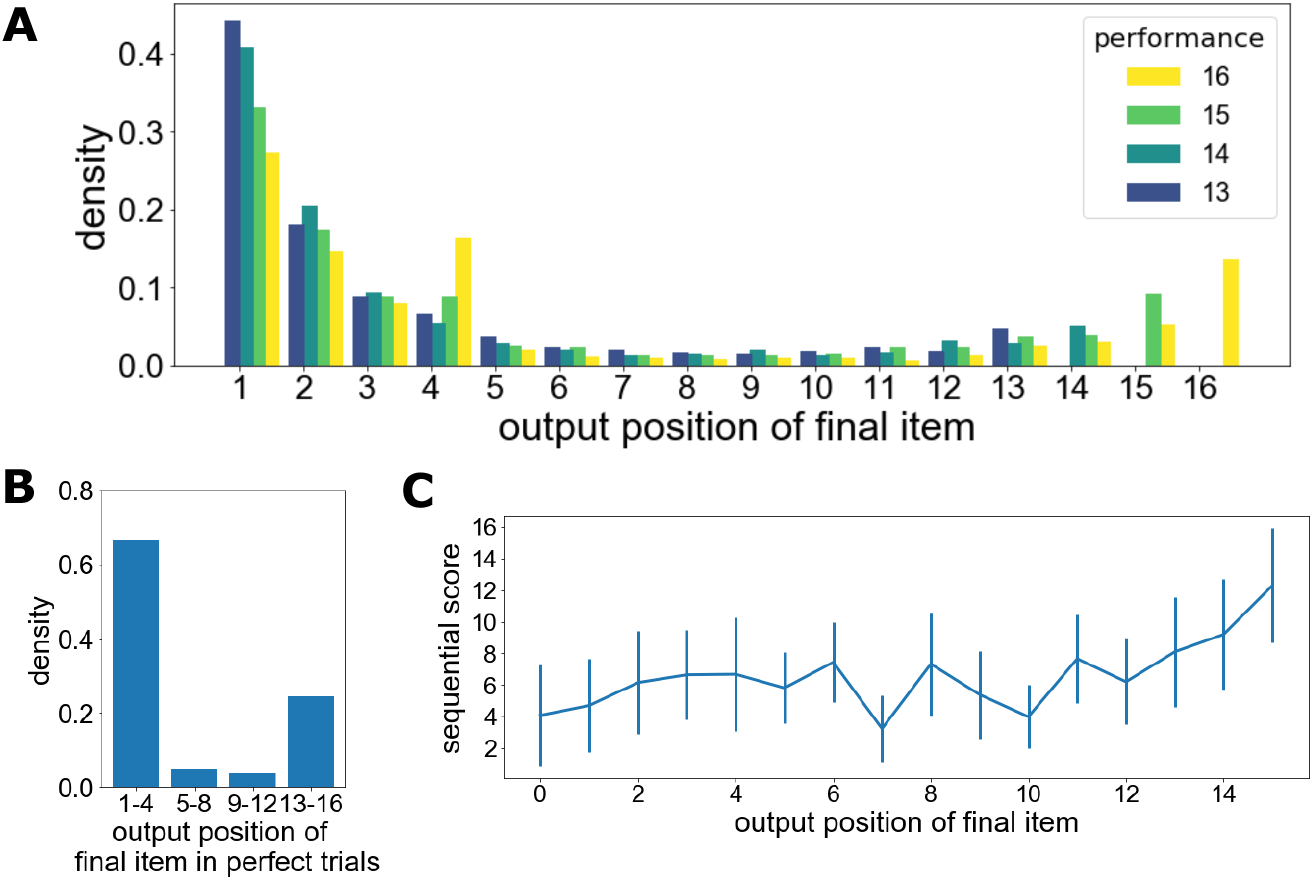
Bimodality of the final item’s output position in high performance trials. **(A)** Full distribution of the final item’s output position over high performance trials. **(B)** Distribution in perfect trials over grouping of four consecutive serial positions. **(C)** Relationships to the sequential score in perfect trials (error bars display standard deviation).

**Figure 15.**
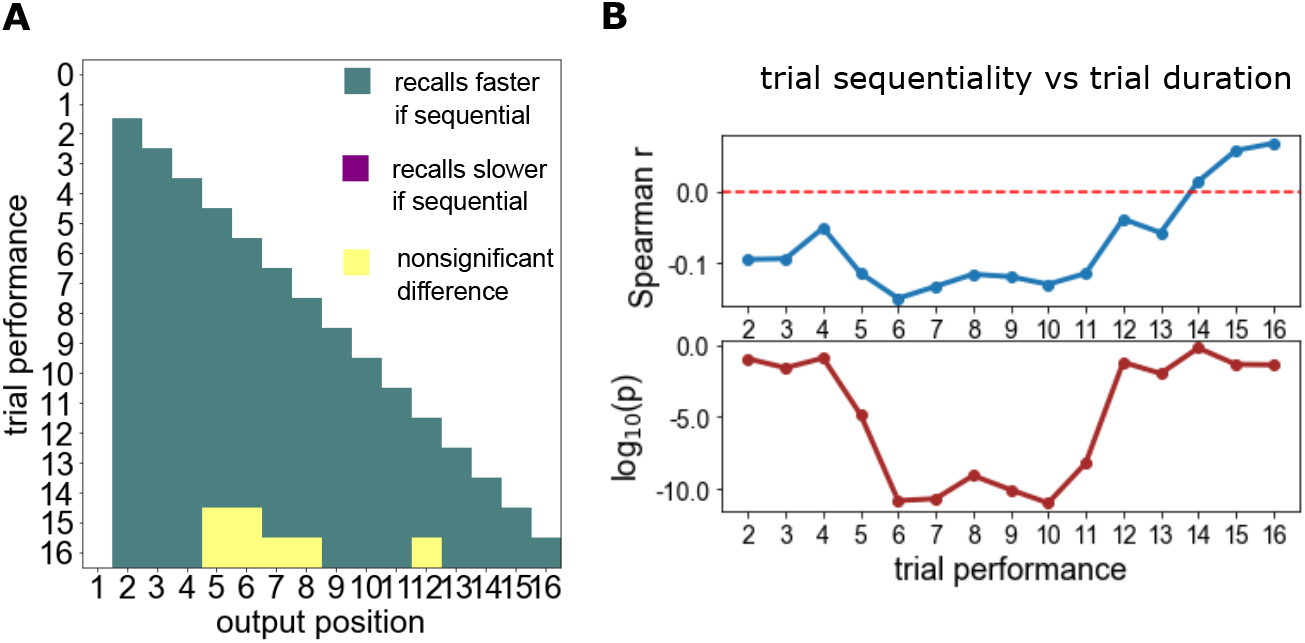
Dependency of recall speed on sequentiality. The dependency is shown here on a recall-by-recall (panel A) and trial-by-trial (panel B) basis. **(A)** Sign of the difference between the mean IRI of sequential and non-sequential recall events, broken down for fixed output positions and trial performances. Color coding: *white* if data are not available from both the sequential and non-sequential conditions; *yellow* if the Mann-Whitney p-value between the two corresponding subsamples is lower than *p*\* = 10^−3^; *green* if sequential recalls are significant faster on average; no case is found where they are slower on average. Spearman’s correlation between overall duration of a trial and its sequentiality, computed over individual performance groups. Significance is found only for mid-performances, with the more sequential trials achieving faster termination.

**Figure 16.**
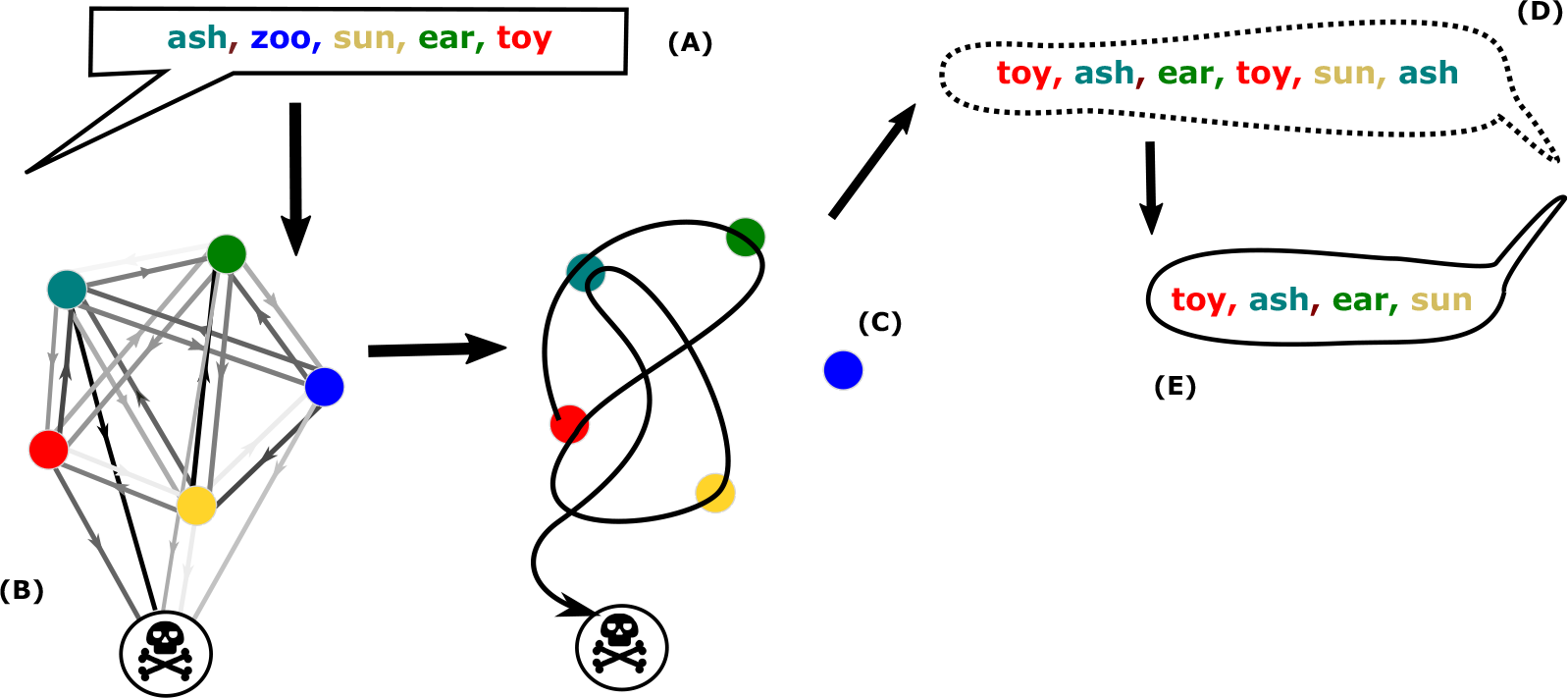
Schematic representation of the skipping model. During the presentation stage **(A)** the subject is presented (visually or orally) with a list of *L* items (in this example, *L* = 5 words). Memories of exposure to each of the items are stored in such a way that mental transitions among those memories have unequal probability, represented in **(B)** by the shade of gray of the arrows. During the recall stage **(C)**, the subject instantiates a mental trajectory through those memories. Nothing prevents the sequence of retrieved memories from containing an unlimited number of repetitions **(D)**, but the output is devoid of repetitions **(E)**, a fact possibly explained by self-censoring.

**Figure 17.**
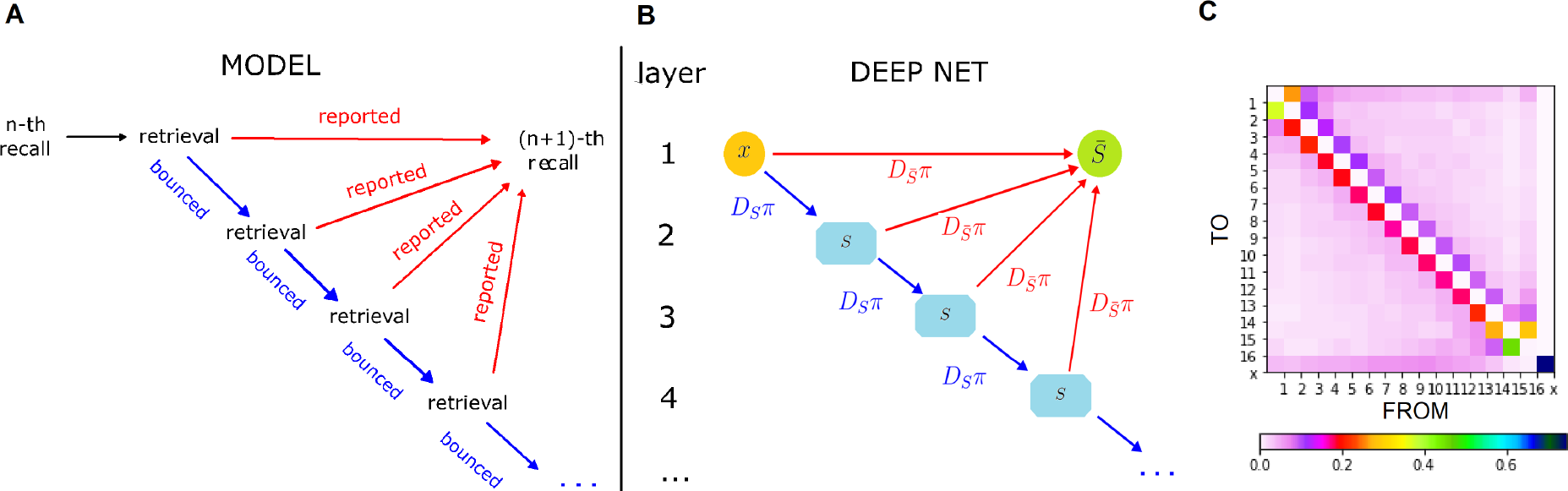
Deep network architecture for fitting the skipping model. The two panels show the correspondence between the“skipping model and its implementation on a deep RNN-style neural network. **(A)** The process occurring between two consecutive recalls is mediated by an arbitrary number of retrieval that go unreported unless they provide a novel item. **(B)** Inter-layer weights in terms of the transition matrix *π* (and in terms of the other matrix operators defined in Supplementary Note 2) represent the operation of bouncing or reporting each retrieval, with each bounce corresponding to a new layer of the network. **(C)** Resulting transition matrix, with the sink state labeled as x.

**Figure 18.**
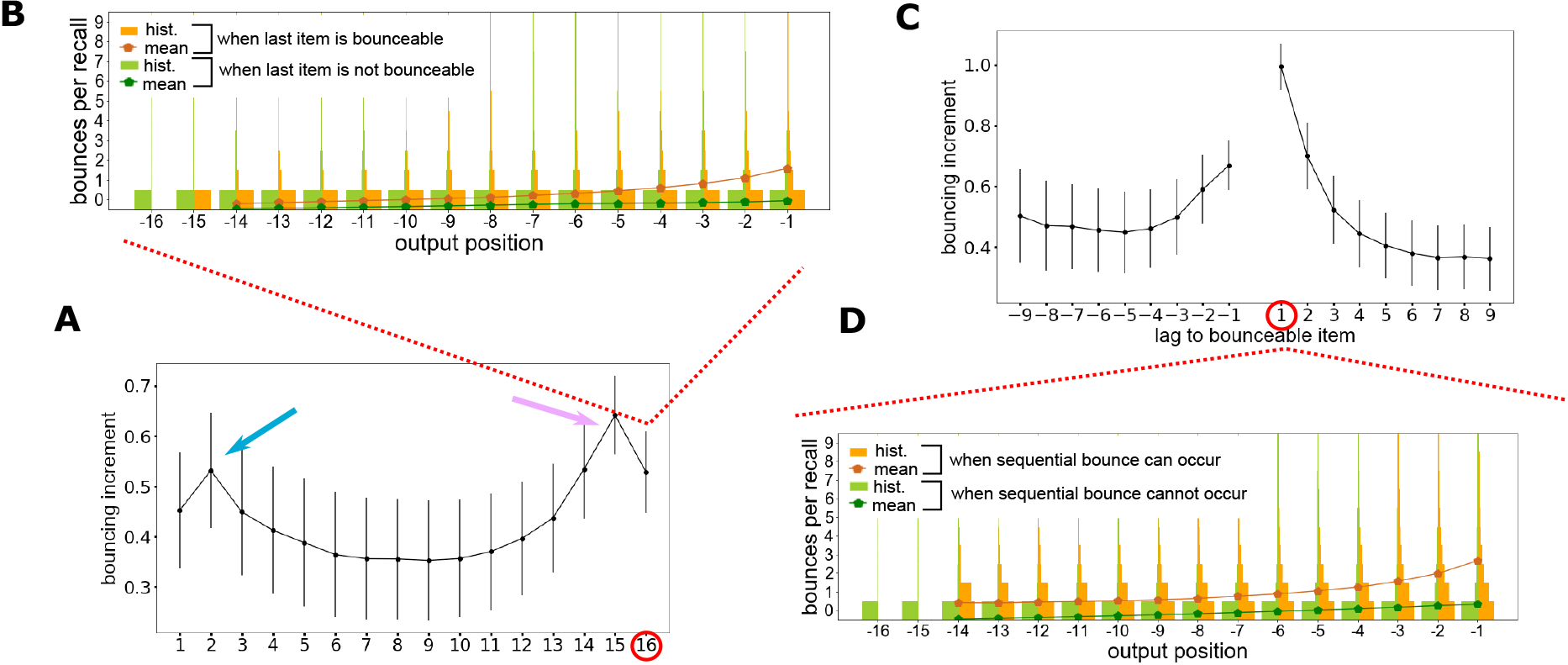
Silent serial-position effects in the skipping model. Panels **(A-B)** correspond, for the skipping model, to panels A-B of Fig. 6 for the bouncing model – and panels **(C-D)** to panels A-B of Fig. 7. Predictions from the skipping model are substantially similar to those of the bouncing model in regard to lag-dependent silent effects. Silent effects depending on serial position depart from bouncing by the stronger emergence of silent primacy (blue arrow), here close to being on the same footing as silent recency.

Notice that sequentiality partially comes from the Markov transition probabilities and partially from strategicity. Our focus is not to achieve the best quantitative model but to explain how the strategicity sequentializes only high performance trials, which yields a major difference between the two mechanisms of increasing sequentiality. For simplicity we used all trials to fit transition probabilities, which in principle may overestimate non-strategic sequentiality.

### Simulations

To run simulations of the bouncing model with and without strategicity, we extract the normalized histogram of the full dataset at output position = 1 and use it as our initial condition. Simulation of 105 trials takes minutes on an ordinary laptop.

Fitting and simulations based on an alternative “skipping model” are described in Supplementary Note 2.

**Table 1.** (Main definitions used in the text)

| **Term** | **Definition** |
| --- | --- |
| Contiguity effect | Tendency to recall contiguous items contiguously |
| Forward asymmetry | Tendency to recall items in forward order |
| Impatience | Probability of quitting the trial per unit bounce |
| Inter-Recall Interval | Time elapsing between two consecutive recalls |
| Lag | Difference between serial positions of the current and latest recall within a trial |
| Negative output position | Number of extant recalls to the end of a trial |
| Output position | Ordered position of an item in the sequence of reported recalls |
| Performance | Number of words explicitly recalled in a trial |
| Primacy effect | Strong recallability of initial items in the list |
| Recency effect | Strong recallability of final items in the list |
| Repetition bouncing | Discarding of a repetitious retrieval |
| Sequential retrieval | A retrieval occurring with lag $L = +1$ |
| Sequential slowdown | Sequential recalls in a given output position being performed slower in more sequential trial |
| Serial position | Ordered position of an item in the presented list |
| Silent contiguity | Recall delay associated to discarded retrievals with lag $L = +1$ |
| Silent recency | Recall delay associated to discarded retrievals of the final item in the list |
| Strategic bouncing | Discarding of non-sequential retrievals to pursue a sequential recall strategy |
| Strategicity | Probability of strategically discarding a non-sequential item if retrieved |
| Trial sequentiality | Fraction of sequential recalls in a given trial |

## Author contributions

The authors worked as a team; credit is not individualizable.

## Availability of data

a used in this work were generously made available by the University of Pennsylvania at the address http://memory.psych.upenn.edu/files/pubs/HealEtal14.data.tgz.

## List of Supplementary Materials

- Supplementary Note 1. Loglikelihood of the bouncing model
- Supplementary Note 2. The “skipping” model
- Table 1. Main definitions used in the text.
- Fig. 10. Overview of recall by serial position of recalled items.
- Fig. 11. Comparison to chance level of the amount of repetitions in the dataset.
- Fig. 12. Terminal Alignment of mean IRIs.
- Fig. 13. Illustration of typical features of the bouncing model.
- Fig. 14. Bimodality of the final item’s output-position in high performance trials.
- Fig. 15. Dependency of recall speed on sequentiality.
- Fig. 16. Schematic representation of the skipping model.
- Fig. 17. Deep network architecture for fitting the skipping model.
- Fig. 18. Silent serial-position effects in the skipping model.

## Supplementary Note 1: Loglikelihood of the bouncing model

In the bouncing model as set up in the main text, upon retrieving a repetitious item, the process reverts to the latest previously retrieved item and makes a new “try” from there. Here we formalize the concept mathematically, writing down the loglikelihood for the model.

Let us label the items by their serial position *x* = 1, …, *N*, where *N* is the length of the memorized list. The first recalled item of every trial is picked according to a distribution *p*_1_(*x*) = *ρ*_init_(*x*) which plays the role of an initial condition. The second item has probability *p*_2_(*x*) = Σ_*x*_1__ *π*(*x*|*x*_1_)*p*_1_(*x*_1_) where *π*(*y*|*x*) is the naked transition matrix. This matrix is assumed diagonal-free, so *π*(*x*_1_|*x*_1_) = 0, and no repetition can occur in the second recall.

At the third recall, for every previous history (*x*_1_*, x*_2_), there is a finite probability *π*(*x*_1_|*x*_2_) of ending up in a repetition. If this happens, the trajectory goes back to *x*_2_ without outputting either retrieval, what we refer to as a “bounce”. In other word there is a set *J* of undesired retrievals that contains in this case only item *x*_1_ (*J* = {*x*_1_}).

If an infinite number of bounces is allowed, the transition from the retrieval of *x*_2_ to the next is governed by a matrix Π_0_(*x*|*x*_2_; *J*) defined as

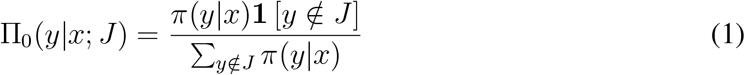

where ***1***[*X*] = 1 if statement *X* is true, 0 otherwise.

In other words, the transition matrix is defined with the target items in the bounceable set masked away, and every other element normalized so the columns sum to unity. Notice the special case Π_0_(*y*|*x*; θ) = *π*(*y*|*x*).

This is however only correct if any number of bounces is allowed; in practice, it is realistic that the subject will give up after a certain number of bounces. Every repetition bounce is thus associated to a fixed surrender probability *q* we call “impatience” (although different modelings of impatience may also be devised).

Call *J_n_* = [*x*_1_, *x*_2_, …, *x*_*n*−1_] the (unordered) set of all items recalled up to time *n*. The probability *p*_*n*+1_(*y*) of item *y* being the *n* + 1-th recall is given by

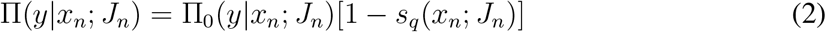

where *s_q_*(*x*; *J*) is the probability of surrendering, i.e. terminating the process, during the recall step starting out from recall of item *x*, given that the bounceable set is *J*.

At each step the process has three possibilities: recalling, bouncing, or surrendering. Recall happens if a new word is retrieved before surrendering; surrendering occurs with probability *q* whenever a repetitious word is retrieved; and bouncing occurs with probability 1 − *q* when a repetitious word is retrieved, returning the process to the latest recalled item.

Calling the three corresponding probabilities *s*, *b*,and *r*, we clearly have

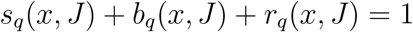

For a single retrieval step, their elemental values are

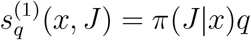

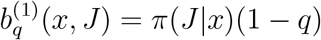

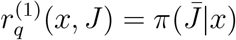

where 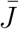 is the allowed set, i.e. the complement of *J*, and we used the notation

*π*(*S*|*x*) = Σ_*y*∈*S*_ π(*y*|*x*) for an arbitrary subset *S* of the *N* list items.

With a potentially infinite number of steps we must write

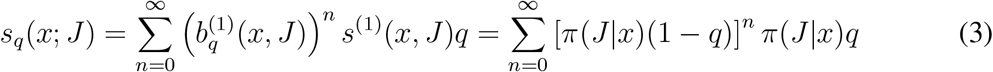

Merging Eqs. (2) and (3), we arrive at

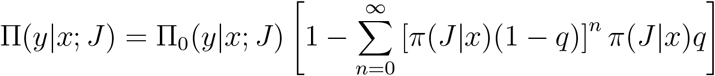

We can now merge in Eq. 1 to eliminate Π_0_, obtaining

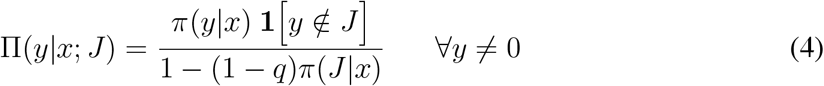

where we made it explicit that this does not provide the probability of transition to silence, which can be described in terms of an effective sink state *y*_sink_ ≡ 0.

The only missing ingredient is now the equivalent of the probabilities (4) for the case of transitions to the sink state. Transitions from the sink state have probability one of staying there and are therefore not problematic:

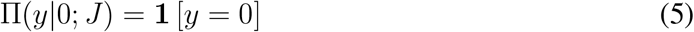

As for the eventuality of recall termination (transition from a list item to the sink), it corresponds to a probability

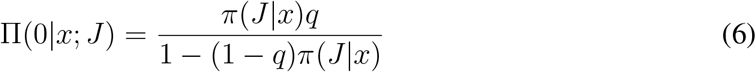

The normalized loglikelihood is a function of the scalar parameter *q* and of the matrix parameter 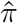 and can be written as

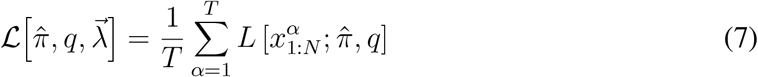

where *T* is the number of trials, and we used the notation

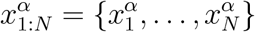

(note that in this convention the last item is also included).

The one-trial loglikelihood appearing in Eq. (7) is defined as

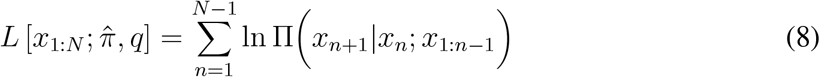

with Π and computed according to Eqs. (4,5,6) for for the given point in the 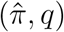 parameter space.

Calling *m_α_* the number of words recalled in the *α*-th trial, we can merge the above formulas into

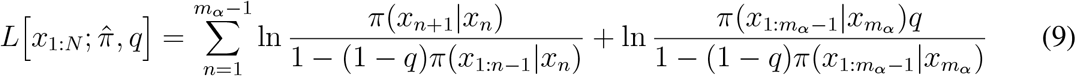

from which, separating the arguments of the logarithms,

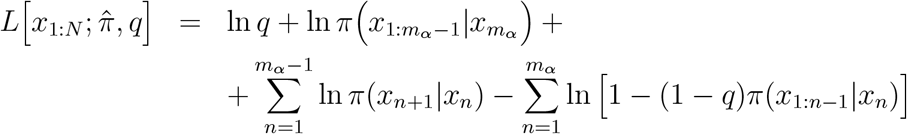

To account for the column-wise normalization of the stochastic matrix Π, instead of using Lagrange multipliers, we opt for adding a degree of freedom by writing

*π*(*y*|*x*) = *u*(*y*|*x*)*/u*(all|*x*) in terms of non-negative auxiliary variables *u*(*y*|*x*) defined for *x* ≠ *y* without any normalization constraint, and with

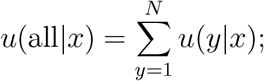

the 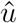-matrix can be assumed to have a zero diagonal.

In terms of the matrix 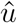, the one-trial loglikelihood (8) becomes

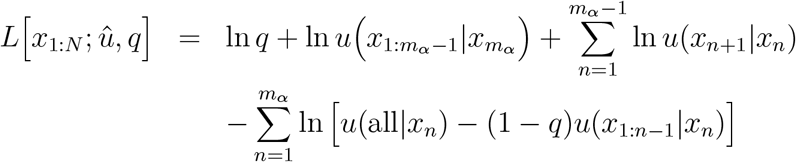

The corresponding q-component of the log-likelihood’s gradient is

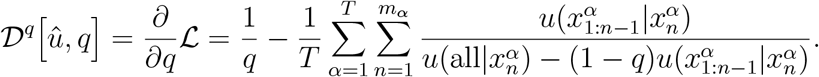

As for the *u*-derivatives, we have

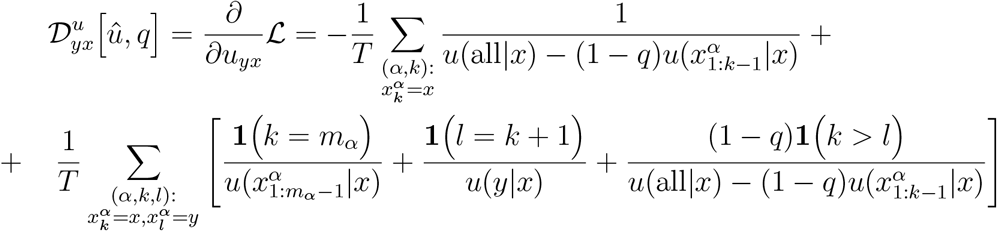

As a sanity check for this last formula, one can isolate the contribution from trials where 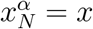 and find that it is exactly zero, as it should because no transitions to the sink are observed from the last recalled item in perfect-recall trials (*m_α_* = *N*), and therefore the likelihood for such trials is not affected by the 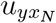 parameter for any *y*.

During the numerical search for an optimum, it is convenient to remove the lower bounds at zero from the elements of 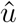 and the [0, 1] bounds from *q*. We do so by mapping both sets of variables to the full real axes. For the sake of convenience we used the transformation

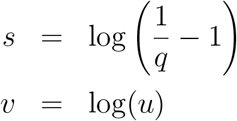

so that the gradient becomes

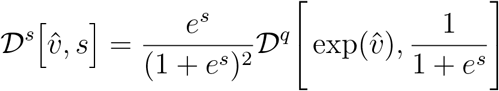

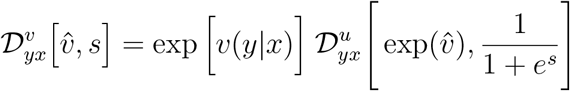

or in terms of the physical variables:

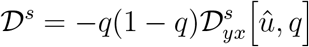

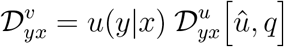

## Supplementary Note 2: The skipping model

An alternative model through which one may attempt to reproduce the silent serial-position effects is what we will refer to as the skipping model, where the markovian retrieval process is not interfered with by the higher-order process that merely censors the reporting of repetitious retrievals.

A repetitious retrieval is thus not discarded as in the bouncing model, but adopted as the starting point for the next transition (what will be described as a “skip” rather than a “bounce”, see Fig. 16).

As in the bouncing model, a *N* × *N* Markov transition matrix *π* is defined such that *π_xy_*is the probability of retrieving word *x* after retrieving word *y*. We assume again that this probability is independent on the time step throughout the retrieval process. We also include among the states of the Markov chain a sink state accounting for termination.

The transition from the *n*-th to the *n* + 1-th recall is governed by a “recall propagator” *T*_{*x*_1_,...,*x_n_*}_, defined as a matrix that depends parametrically on the potentially repetitious (hence unreportable) set of serial positions {*x*_1_*, …, x_n_*}, which is the set of words recalled up to that moment.

In other words, for any set of serial positions *S*, we are defining the matrix *T_S_* such that [*T_S_*]*_xy_* is the probability of recalling *x* after recalling *y*, given that the items in the set *S* cannot be reported having already been recalled (though they can be retrieved). Since recalls do not happen at all retrieval events, skipping repetitious retrievals, for *x* ∈ *S* or *y* ∉ *S* we have [*T_S_*]*_xy_* = 0, (Fig. 17A).

Since the dataset samples transitions driven by the *T_S_* matrix, the *π* matrix can be obtained via maximizing the likelihood of *T_S_* Matrix given the dataset. Unfortunately, even if infinite skips are allowed in a single recall event and no sink state included, a closed formula like Eq. (1) is not available for *T_S_*. Although a Dyson-like summation can be performed for this propagator, when computing the loglikelihood on data implementing it leads to a combinatorial blow-up of the number of possible paths.

Thus, while the retrieval process in the skipping model is simpler than in the bouncing model (because it stays completely Markovian) the fitting is much less straightforward.

On the other hand, once the *π* matrix is given, the *T* matrix can always be calculated iteratively. In order to practically estimate the *π* matrix, we exploited the fact by making an extra assumption and putting an upper limit *n*_max_ on the possible number of bounces. We were thus able to build an RNN-style computational graph as illustrated in Fig. 17B.

For any set of serial positions *S*, let us define the matrix *D_S_* such that

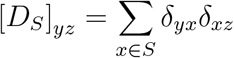

Given a certain source item *x* and a repetitious set *S + x = S* ∪ *x*, the *T* matrix can be calculated as:

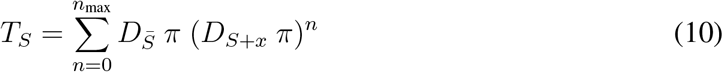

where 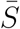 is the allowed output set and *S* + x is the unreportable (i.e. reptitious) set; *x*, which is the source, is included in the unreportable set.

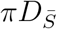 is here a skip-connect indicating end of a bounce when an element recalled is in the reportable set. This graph now also allows transitions from *x* to itself, a hitch that can be solved by masking out diagonal elements of *π* matrix. Even without masking, however, the diagonal elements of *π* matrix vanishes after fitting the data.

We implement the computational graph with PyTorch. Negative log-likelihood loss function between calculated source-to-target log-probability with current *π* Matrix and observed target sample is used as the loss function. A certain batch number of unreportable sets and reported targets is randomly picked form the data set to carry out a stochastic-gradient-descent parameter update of the *π* Matrix.

Let us call 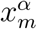 the serial position of the word recalled in the m-th recall of trial *α*. Call *b* the size of minibatches. Each minibatch selection consists of a set 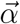 of *b* random trial indices *k* and of a set 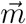 of corresponding random output positions *m_k_*. Here *k* = 1*, …, b*, and for each *k*, 1 ≤ *k* ≥ *N_T_* (*N_T_* being the number of trials) and 1 ≤ *m_k_* ≥ *L* (L being the list length).

The loss function for a particular batch is then

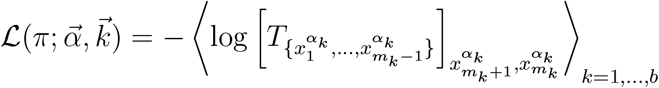

where average is used instead of sum to keep the values of the loss function numerically low even with many batches. It takes approximately tens of minutes to obtain convergence on a conventional laptop.

The resulting matrix parameter *π*, shown in Fig. 17C can be used to simulate the model starting from the empirically observed initial condition. Both the silent recency effect and the silent contiguity effect are successfully reproduced (Fig. 18).

However, an additional “silent primacy effect” (where past recall of the first item in the list engenders a delay in subsequent recalls) emerges nearly as conspicuously as silent recency. This is compatible with the existence of a corresponding primacy effect in reported recall frequencies (Murdock Jr, 1962) but is not compatible with the data (Fig. 3).

